# Microthrombi-growth in ADAMTS13 deficiency exacerbated ulcerative colitis via mucosal and endothelial dysfunction

**DOI:** 10.1101/2024.12.10.627861

**Authors:** Kyota Tatsuta, Naoki Honkura, Nanami Morooka, Mayu Sakata, Kiyotaka Kurachi, Ken Sugimoto, Koichi Kokame, Hiroya Takeuchi, Tetsumei Urano, Yuko Suzuki

## Abstract

**Background:** Ulcerative colitis (UC) is a chronic inflammatory bowel disease characterized by mucosal inflammation and ulceration, with systemic immune dysregulation exacerbating disease progression. While the accumulation of von Willebrand factor (VWF), normally digested by a disintegrin-like and metalloproteinase with thrombospondin type 1 motif 13 (ADAMTS13) from ultra-large multimers to small molecules, has been implicated in UC pathogenesis, the association between aberrant microthrombus formation and colitis remains unclear.

**Methods:** Plasma and inflamed colonic tissues from UC patients were analyzed. In the experimental colitis model, we employed an intravital imaging technique to reveal real-time structural dynamics of the mucus layer and the mucosal vasculature. Dextran sulfate sodium (DSS)-induced colitis in wild-type and ADAMTS13-deficient mice was evaluated for disease progression, mucus layer disruption, leukocyte recruitment, and thrombus formation in mucosal vessels using intravital multiphoton excitation microscopy within single-cell spatial resolution.

**Results:** UC patients exhibited significantly reduced plasma ADAMTS13 activity correlating with disease severity and excessive VWF deposition in inflamed colonic tissues. In DSS-induced colitis mice, ADAMTS13 deficiency showed heightened disease activity and increased mucosal erosion in histochemical analysis compared to wild-type mice. A novel methodology established in this study using intravital microscopy successfully appraised colonic mucus barrier integrity by visualizing fluorescent dextran penetration from the colonic lumen to the crypts. ADAMTS13 deficiency accelerated mucus layer disruption, leukocyte adhesion, and microthrombi formation, particularly close to crypt epithelium regions. Vessel-specific analyses demonstrated that obstructive microthrombi were most prominent in the mucosal layer, contributing to local ischemia and mucosal erosion. Therapeutical usage of recombinant human ADAMTS13 alleviated microthrombus formation, improved mucosal integrity, and mitigated colitis severity in wild-type and ADAMTS13-deficient mice.

**Conclusions:** Obstructive thrombi formed in mucosal vessels due to impaired ADAMTS13 activity appeared essential in the disease progression of UC. Advanced intravital imaging provided novel insights into single-cell resolution UC pathogenesis.

## Introduction

Ulcerative colitis (UC) is an inflammatory bowel disease that is characterized by inflammation and mucosal ulceration that extends from the rectum to the colon.^1^ The early stages of pathogenesis in UC involve disruption of the mucus layer, which comprises goblet cell secretory components, consequently the loss of mucosal homeostasis.^2^ Subsequently, the dysregulated activation of the mucosal immune system exacerbates colonic mucosal inflammation and leads to the progression of systemic inflammation.^3^ Despite significant advancements in therapeutics through the utilization of molecularly targeted drugs that are based on immune signals,^4^ clinical trials continue to demonstrate low remission rates.^5,6^ A comprehensive understanding of the intricate pathophysiological mechanisms that both initiate and exacerbate UC, including mucus layer disruptions and inflammatory responses within the mucosal microvasculature, is crucial for enhancing disease management strategies.^7^

In these responses, von Willebrand factor (VWF) has emerged as a significant factor, closely associated with endothelial dysfunction in UC.^8^ A disintegrin-like and metalloproteinase with thrombospondin type 1 motif 13 (ADAMTS13) cleaves ultra-large (UL)-VWF multimers, which are stored in the Weibel-Palade bodies of endothelial cells and released into the blood in response to various stimuli, into smaller and less reactive fragments with blood flow.^9^ Inflammatory diseases inhibit ADAMTS13 synthesis via inflammatory cytokines (e.g., interleukin-6 and tumor necrosis factor-α) and inactivate ADAMTS13 through proteases released from neutrophils during inflammation,^10^ resulting in the inability to cleave UL-VWF multimers and potentially triggering thrombus formation. Clinical studies have suggested that decreased ADAMTS13 activity is associated with an increased risk of thromboembolic events in patients with UC.^11^

A previous study using UC model mice and human tissue sections revealed that ADAMTS13 deficiency led to VWF-rich thrombus formation in submucosal vessels, then exacerbating colitis.^12^ The administration of recombinant human ADAMTS13 (rhADAMTS13) reduced the severity of colitis, suggesting its potential as an adjunct to existing therapies.^12^ The presence of thrombi in mucosal microvessels has been reported in several studies analyzing rectal biopsies of patients with UC.^13,14^ However, these findings remain inconclusive, as pathological analyses were conducted exclusively on colon tissue sections. Consequently, the causal relationship between microvessel thrombi and disease severity has yet to be established. Accurately identifying the spatial and temporal distribution of thrombi within mucosal vessels during UC progression significantly enhances the understanding of the underlying pathophysiological mechanisms.

In decades, advances in real-time imaging techniques have enabled not only in vitro analyses ^15–17^ but also the study of tissue in living animals,^18–21^ allowing us to assess the spatiotemporal dynamics in single platelets and vascular endothelial cells. We investigated the spatiotemporal processes of platelet activation, coagulation, and fibrinolysis by employing a laser-induced model of single endothelial cell injury in mice.^19–21^ Advanced live imaging techniques utilizing multiphoton excitation microscopy were further developed to achieve high spatial resolution, thereby enabling the observation of individual cell dynamics.^22^

In this study, we employed low-invasive intravital imaging with multiphoton excitation microscopy to investigate colonic lesions in colitis mice and attempted to elucidate the spatiotemporal regulatory mechanisms that exacerbate colitis. We propose potential linkages among mucus layer dysfunction, the increased formation of obstructive thrombi in the mucosal microvessels near the epithelial layer, and the progression of disease, utilizing an ADAMTS13-deficient microthrombosis model in mice.

## Methods

Details can be found in the Supplemental Materials.

### Human sample preparations and analysis

Ethical approval for using human samples was obtained from the IRB of Hamamatsu University School of Medicine (approval no. 21-208). We obtained plasma and measured plasma ADAMTS13 activity in samples from patients with UC and healthy volunteers. Immunohistochemical analysis of VWF staining was conducted to compare the most inflamed regions of UC tissues with the normal colon, defined in this study as non-cancerous areas from Stage 0 or I colorectal cancer specimens.

### Animals

The green fluorescent protein (GFP)-expressing transgenic C57BL/6J mice were supplied by Dr. Okab.^23^ *Adamts13*-deficient mice (C57BL/6 strain background) with confirmed activity deficiency due to the lack of cleavage of GST-mVWF73-H and FRETS-VWF73, which are specific substrates of ADAMTS13,^24^ were crossed with GFP mice (ADAMTS13-KO mice).^21^ All experiment animals were 7-to 8-week-old males and weighed between 20 and 23 g. Age-stratified data on body weight, autoclaved water consumption, and food intake are shown in Figure S1. After the completion of the experiments, all mice were euthanized to minimize unnecessary suffering.

### Dextran sulfate sodium-induced colitis mouse model

Acute colitis was induced by the oral administration of 5% (w/v). DSS (MP Biomedicals) in autoclaved water for 5 days. Daily weight and clinical colitis scores were measured using the disease activity index^25^ based on weight loss (0: none, 1: 1–5%, 2: 5–10%, 3: 10–15%, and 4: >15%), stool consistency (0: normal, 2: loose stool, 4: watery diarrhea). and the degree of intestinal bleeding (0, normal; 2, slight bleeding; and 4, gross bleeding).

### Histological assessment in the DSS-induced colitis mouse

We evaluated inflammation-associated histological changes in the colon, the number and length of the mucosal erosions, and the number of periodic acid Schiff (PAS)-positive goblet cells per colonic crypt.

### Intravital imaging system

Intravital imaging was conducted using a Nikon A1R MP multiphoton microscope (Nikon) equipped with an ultrashort-pulse tunable Ti:sapphire laser (Chameleon Vision II, Coherent). For two-photon excitation, wavelengths were set to either 960 nm or 1040 nm, tailored to each experimental condition. Emitted fluorescence signals were split sequentially into green and red channels via dichroic mirrors (560 nm and/or 593 nm), optimizing for each excitation wavelength and fluorescence molecule brightness to prevent cross-talk. The fluorescence was then filtered through band-pass emission filters (525/50 nm for green [GFP] and 629/56 nm for red [TRITC: Tetramethylrhodamine or Alexa Fluor 568]) and captured separately using GaAsP photomultiplier tubes.

### Recombinant human ADAMTS13 treatment

A single shot of 0.5 µg RhADAMTS13 (R&D Systems) was administered by retro-orbital intravenous injection every 24 h on days 1–5 after DSS administration. All experiments were performed four hours after rhADAMTS13 administration.

## Results

### Impaired ADAMTS13-VWF axis correlates with disease severity in UC

First, we evaluated the effect of the ADAMTS13-VWF axis in patients with moderate-to-severe UC, with disease severity defined according to the Endoscopic Mayo Score (Supplemental methods for detailed classification).^26^ In blood samples collected from patients with moderate-to-severe UC before administering inpatient treatment, plasma ADAMTS13 activity levels were significantly lower than in healthy controls (Figure 1A). Severe cases of UC showed weaker ADAMTS13 activity than moderate cases, although the difference was not significant. We undertook a detailed evaluation of the correlation between plasma ADAMTS13 activity and plasma albumin levels, which are negatively correlated with disease progression, and platelet counts, which are positively correlated with disease progression,^27^ and found the correlation of ADAMTS13 activities similar to the disease progression (Figure 1B and 1C). To clarify how decreased ADAMTS13 activity affects VWF distribution in lesions, we performed VWF staining in sections of inflammatory colon obtained from patients with UC. In normal colon tissue sections, anti-VWF immunostaining showed VWF deposition only in the vessel walls. In contrast, the inflamed tissue of patients with UC showed significant VWF accumulation in the vessel walls and interstitial spaces of the mucosal and submucosal layers (Figure 1D and 1E). We quantitatively analyzed the percentage of VWF-stained areas in the mucosal and submucosal layers across the entire colonic tissue section. A significant increase in VWF accumulation was observed in the inflamed UC colon compared to the normal colon (Figure 1E). In addition, the amount of *VWF* mRNA in the colonic tissue of patients with active UC was higher than that in the normal colon (Figure S2), which was analyzed using two different datasets.^28,29^ These results indicate that UC disease severity could be characterized by the active levels of ADAMTS13 and VWF in inflamed lesions.

**Figure 1.**
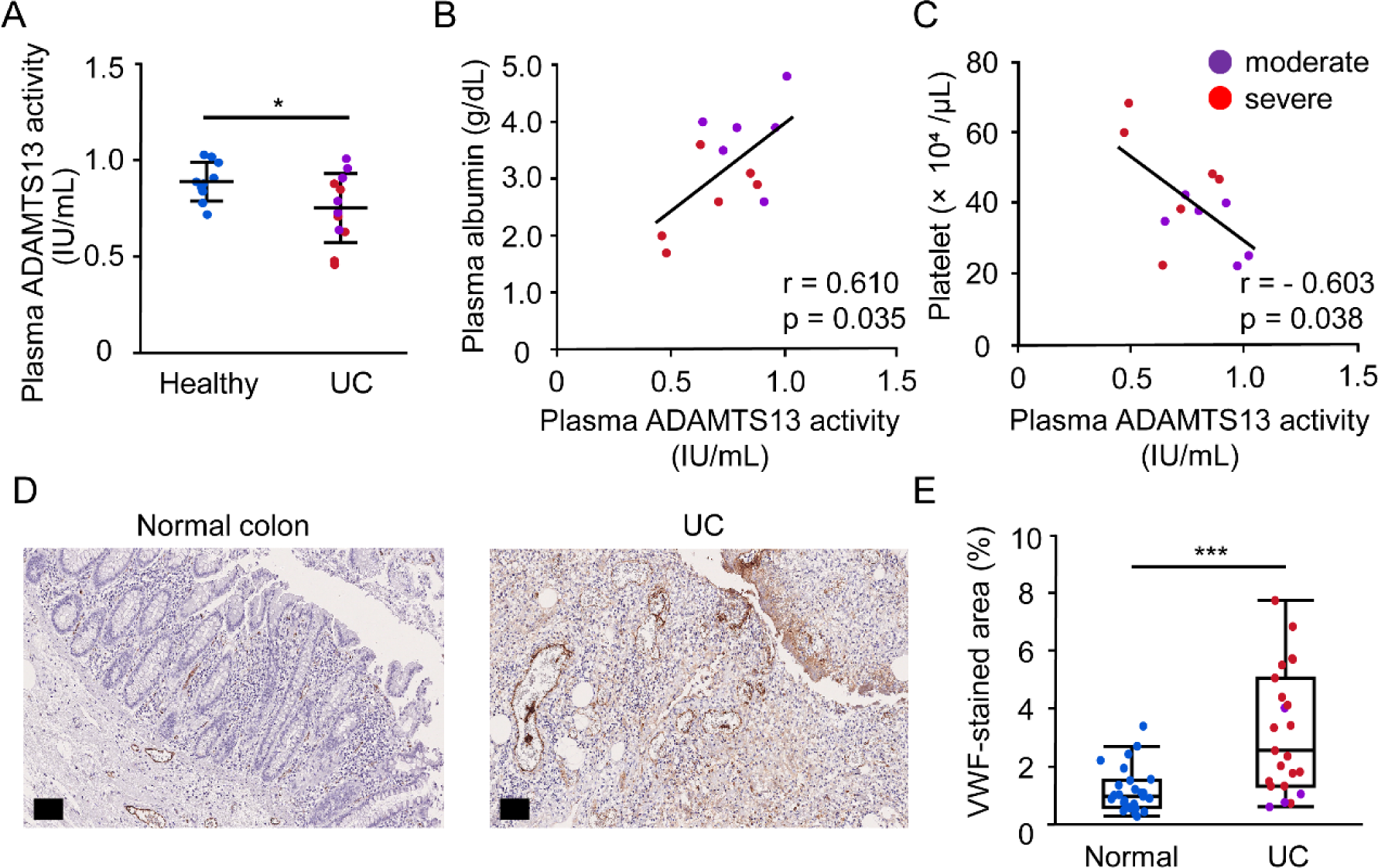
Decreased ADAMTS13 activity and increased VWF level in patients with UC. (A) Comparison of plasma ADAMTS13 activity in healthy controls (n=10, blue dots) and patients with UC (n=12). UC disease severity was defined as moderate and severe based on Endoscopic Mayo Scores of 2 and 3 (n=6 each, purple and red dots), respectively. (B, C) The relationship of plasma ADAMTS13 activity levels with plasma albumin levels and platelet counts in UC. (D) Representative images of VWF immunohistochemistry in colonic tissue sections from normal colon and those with UC; scale bar: 100 µm. (E) Quantitative analysis of VWF-stained areas in colonic surgical specimens from normal (n=27) and inflamed UC (n=24) participants. Data (A), presented as mean and SD, were analyzed by the Student’s t-test. Data (B, C) were analyzed by the two-sided Spearman’s correlation. Data (E), presented as median (center line) with 25th and 75th percentiles (box bounds) and whiskers (maximum and minimum data points), were analyzed by the Mann–Whitney U test. *p<0.05; ***p<0.001.

### ADAMTS13 deficiency exacerbates DSS-induced colitis in mice

To understand the pathophysiological mechanism of colitis exacerbated by reduced ADAMTS13 activity, we induced colitis in ADAMTS13-KO mice using DSS, which shows many similarities with human UC.^30^ Oral administration of 5% DSS induced similar weight loss in WT and ADAMTS13-KO mice (Figure 2A), whereas the disease activity index, as evaluated by weight loss, stool consistency, and the degree of intestinal bleeding, was more severe in ADAMTS13-KO mice than in WT mice (Figure 2B). The high disease activity index was mainly influenced by increased intestinal bleeding. After 5 days of DSS administration, ADAMTS13-KO mice tended to show shorter colon lengths, a characteristic of DSS-induced colitis in mice,^31^ compared to WT mice (Figure 2C). Mucosal erosion in pathological colon sections was more common in the ADAMTS13-KO mice with DSS-induced colitis than in the WT mice (Figure 2D, black arrowheads). Histopathological scores, evaluated by the degree of tissue damage and inflammatory cell infiltration,^31^ were significantly higher in ADAMTS13-KO mice with DSS-induced colitis (Figure 2E). The total number of mucosal erosions (Figure 2F) and the normalized length of mucosal erosion (Figure 2G) in the entire colon were significantly increased in ADAMTS13-KO mice than in WT mice. Our findings indicate that ADAMTS13 deficiency significantly exacerbates DSS-induced colitis, particularly at sites of mucosal erosion in mice.

**Figure 2.**
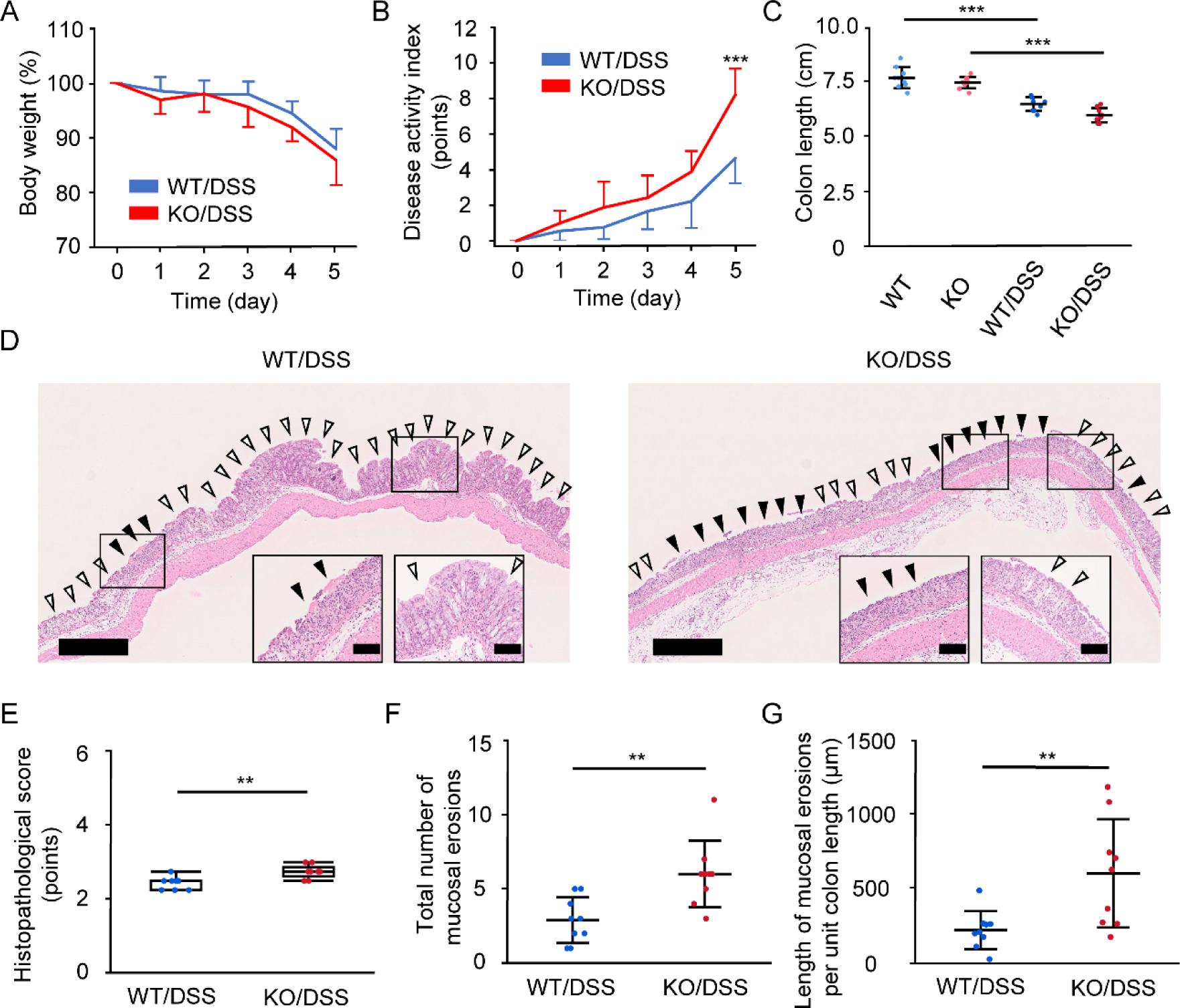
The impact of ADAMTS13 deficiency on DSS-induced colitis in mice. Mice (age: 7–8 weeks, male, n=9 per group) were administered 5% DSS for 5 days. (A, B) Changes in body weight and the disease activity index from 0 to 5 days (blue lines, WT mice; red lines, ADAMTS13-KO mice). (A) Body-weight changes are shown as percentages of those before DSS administration (day 0). (B) Disease activity indices (0, no symptoms, to 12, most severe) were calculated daily based on weight loss, stool consistency, and degree of intestinal bleeding. (C) The total colon length from the proximal colon to the rectum was measured on days 0 and 5 (light blue dots, WT mice on day 0; light red dots, ADAMTS13-KO mice on day 0; blue dots, WT mice on day 5; red dots, ADAMTS13-KO mice on day 5). (D) Representative images of H&E staining of DSS-induced inflamed colon sections of WT and ADAMTS13-KO mice on day 5 (black and white arrowheads: areas of mucosal erosion and where the surface epithelium is maintained, respectively; scale bars: 500 and 100 µm [inset]). (E) Histopathological scores (0, no inflammation, to 6, most severe inflammation) on day 5. (F) Number of mucosal erosions in the entire colon on day 5. (G) The length of mucosal erosions per 1 cm total colon length on day 5. Data presented as (A–C) mean and SD, analyzed by the Student’s t-test with Bonferroni correction, (E) Median (center line) with 25th and 75th percentiles (box bounds) and whiskers (maximum and minimum data points), analyzed using the Mann–Whitney U test, and (F, G) mean and SD, analyzed using the Student’s t-test. **p<0.01; ***p<0.001.

### Intravital imaging analysis of mucus layer barrier function

To investigate the early-phase dynamics of UC pathophysiology, particularly concerning the barrier function of the mucus layer (<1-μm thickness), we employed intravital imaging using multiphoton excitation microscopy with fluorescent molecules. The colonic mucus layer comprises a dual-layer system: a densely packed inner layer firmly attached to the epithelium and a fluidic outer layer serving as a protective filter that physically separates bacteria from epithelial cells.^32^ By administering fluorescent dyes into the colon lumen, we attempted to visualize the barrier function of the mucus layer, wherein the fluid mucus dynamics are usually undeterminable with traditional biological methods. TRITC-dextran (40 kDa), a typical biomolecular weight, was injected into the lumen to evaluate its distribution across the colonic crypts, highlighting changes as the mucus layer progressively degraded under UC conditions. After 15 minutes, the colonic lumen was adequately filled with TRITC-dextran, and images of the mouse’s proximal colon were captured by fixing it with agarose onto a glass slide (Figure 3A). TRITC-dextran (red) penetrated the lower parts of the crypts in WT mice with colitis on day 5 post-DSS administration, contrasting with WT mice without colitis, where no such penetration was observed (Figure 3B; Figure S3A; video S1 and S2). To quantify TRITC-dextran distribution, regions of interest (ROIs) were set within the colonic crypts (Figure S3A), and the red signal to the background (green channel) ratio was determined to account for inter-individual variability. The stable GFP fluorescence originated from cells that drove a modified chicken β-actin promoter in the GFP mice.^23^ Notably, cells of the same type exhibited highly consistent GFP expression levels across different individuals. This consistency allowed us to use green fluorescence as a constant reference, enabling precise determination of the red-to-green intensity ratio. By normalizing the red intensity to this stable green fluorescence, we corrected for variability due to interindividual differences among mice and minor imaging fluctuations, such as laser power variations along the z-axis (Figure 3B).

**Figure 3.**
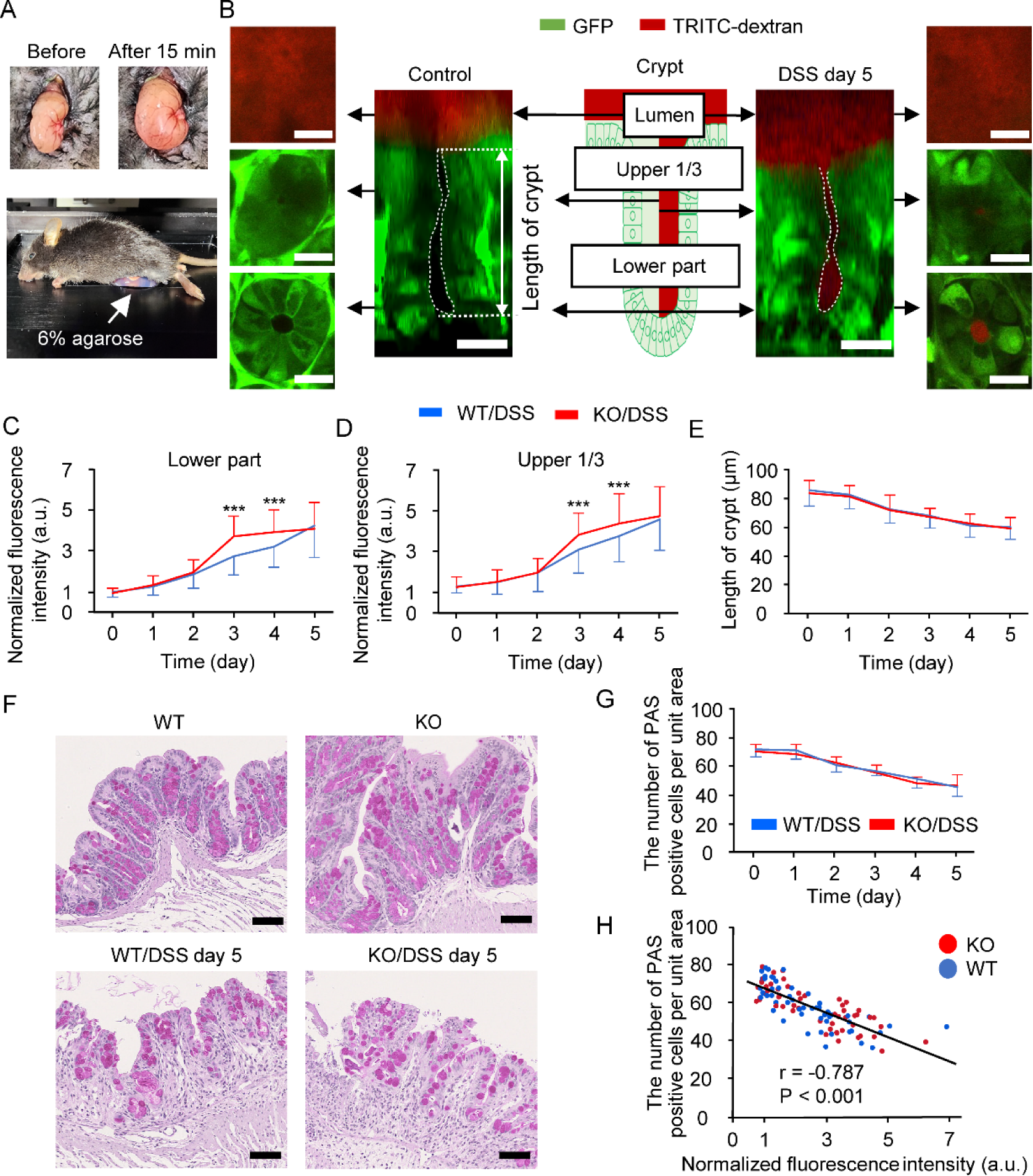
Functional evaluation of the colonic mucus layer. Mice (age: 7–8 weeks, male, n=9 per group) were administered 5% DSS for 5 days. (A) Proximal colon (fixed on 6% agarose) before and 15 min after dextran administration. (B) TRITC-dextran (red) distribution of colonic crypts in the control and DSS-induced colitis groups (day 5; scale bar: 20 µm). (C, D) Normalized fluorescence intensity of TRITC-dextran in the lower (C) and upper one-third (D) crypts. (E) Colonic crypt length (measured at the site indicated in B). (F) Representative images of periodic acid-Schiff (PAS) staining of proximal colon sections prepared from the intravital imaging area (goblet cells were stained purple; scale bar: 50 µm). (G) The number of PAS-positive cells per 100 µm2 of the crypts in DSS-induced WT (blue) and ADAMTS13-KO (red) mice. (H) Correlation between the normalized fluorescence intensity of TRITC-dextran in the lower part of the crypts and the number of goblet cells in the intravital imaging area. Data (mean ± SD) were analyzed by the Student’s t-test with Bonferroni correction (C–E and G) and the two-sided Spearman’s correlation (H). ***p<0.001.

### Impact of ADAMTS13 deficiency on fluidic mucus layer barrier function

The TRITC-dextran fluorescence baseline ratio (day 0) was established for WT mice pre-DSS administration (Figure S3C). The DSS exposure increased fluorescence at both the lower and upper crypt regions in WT and ADAMTS13-KO mice, indicating barrier permeability changes (Figure S3B and S3C). Across all observed points, fluorescence in the upper crypt consistently exceeded that of the lower part, regardless of genotypes (Figure 3C and 3D; Figure S3D). Increased normalized fluorescence intensity on days 3 and 4 in ADAMTS13-KO mice, alongside enhanced disease activity and histological damage, suggested a deficiency-associated reduction in fluidic mucus layer integrity. No significant crypt length differences were observed between the two genotypes (Figure 3E). Together, these findings indicate a DSS-induced expansion of TRITC-dextran distribution from the lumen toward the lower crypt part due to diminished mucus, implicating ADAMTS13 deficiency in mucus layer reduction and endothelial dysfunction at early stages post-DSS exposure.

### Reduction of fluidic mucus layer and goblet cells following DSS administration

We further assessed the correlation between TRITC-dextran penetration and goblet cell depletion in the proximal colon. Reduced mucus secretion is known to correlate with a decrease in goblet cell count.^33^ To assess this, we evaluated goblet cells in the proximal colon using intravital imaging, excluding regions of mucosal erosion. The number of goblet cells (PAS-positive cells) within colonic crypts was determined by averaging counts from 10 crypts and normalizing them to a surface area of 100 µm² per crypt. DSS administration resulted in a time-dependent reduction of goblet cells in both genotypes (Figure 3F and 3G; Figure S4A). However, no significant differences in goblet cell numbers were noted between genotypes on days 3 and 4 post-DSS administration (Figure 3G). These observations suggest that DSS-induced damage to goblet cells, accompanied by endothelial dysfunction, may impair mucus release, impacting both cell counts and mucus volumes in colonic crypts. This assessment, however, presents technical challenges without our novel TRITC-dextran-based methodology (Figure 3C, 3D, and 3G; Figure S4A). A negative correlation was found between normalized fluorescence intensity at the crypt’s lower and upper and goblet cell count, which correlated with DSS exposure duration (Figure 3H; Figure S4B and S4C).

### ADAMTS13 deficiency enhances leukocyte recruitment in DSS-induced colitis

Disruption of the intestinal barrier promotes inflammatory cytokine production and immune cell recruitment to the bowel wall.^34^ We continuously examined early-stage events in DSS-induced colitis, focusing on the subepithelial region and mucosal microvessels. To elucidate the spatiotemporal remodeling of mucosal microvessels through inflammation, we observed leukocyte rolling and adhesion in the microcirculation using high-spatiotemporal resolution intravital imaging. Leukocytes, identified as GFP-positive cells (∼10 µm, white arrows in Figure S5A and S5B; video S3 through S6), were tracked in the circulation. From day 4 of DSS treatment, WT mice displayed increased leukocyte rolling in both small vessels (10–50 µm, Figure S6A) and larger vessels (>50 µm, Figure S6B). Notably, both genotypes of mice exhibited enhanced leukocyte rolling and adhesion to the endothelial wall on day 5, but this increase emerged earlier, from day 3, in ADAMTS13-KO mice (Figure S5C through S5F; Figure S6). These results suggest that ADAMTS13 deficiency accelerates leukocyte recruitment resulting from UL-VWF multimer-related endothelial response, then impacts microcirculatory changes in the inflamed intestine during diminished mucus integrity.

### ADAMTS13 deficiency promotes microthrombus formation within mucosal and submucosal vessels in DSS-induced colitis

Activated endothelial cells by inflammation in the intestinal microvasculature, alongside immune responses, could play a critical role in the early stages of UC. We investigated the impact of endothelial cell-released VWF and the presence or absence of ADAMTS13 on platelet adhesion and microthrombus formation within DSS-induced inflammatory lesions in the mucosal and submucosal vessels. By using Alexa Fluor-568-labeled anti-VWF antibodies, we imaged VWF within inflamed vessels and observed platelet adhesion to one or two layers of VWF multimers (width < 10 µm) and microthrombi formed by aggregated platelets (width ≥ 10 µm) (Figure 4A), similar to the images reported in the previous study.^21^ Grossly, on day 5 post-DSS administration, microthrombi were larger and more prominent in ADAMTS13-KO mice than in WT mice (Figure 4B, white arrows; video S7 and S8).

**Figure 4.**
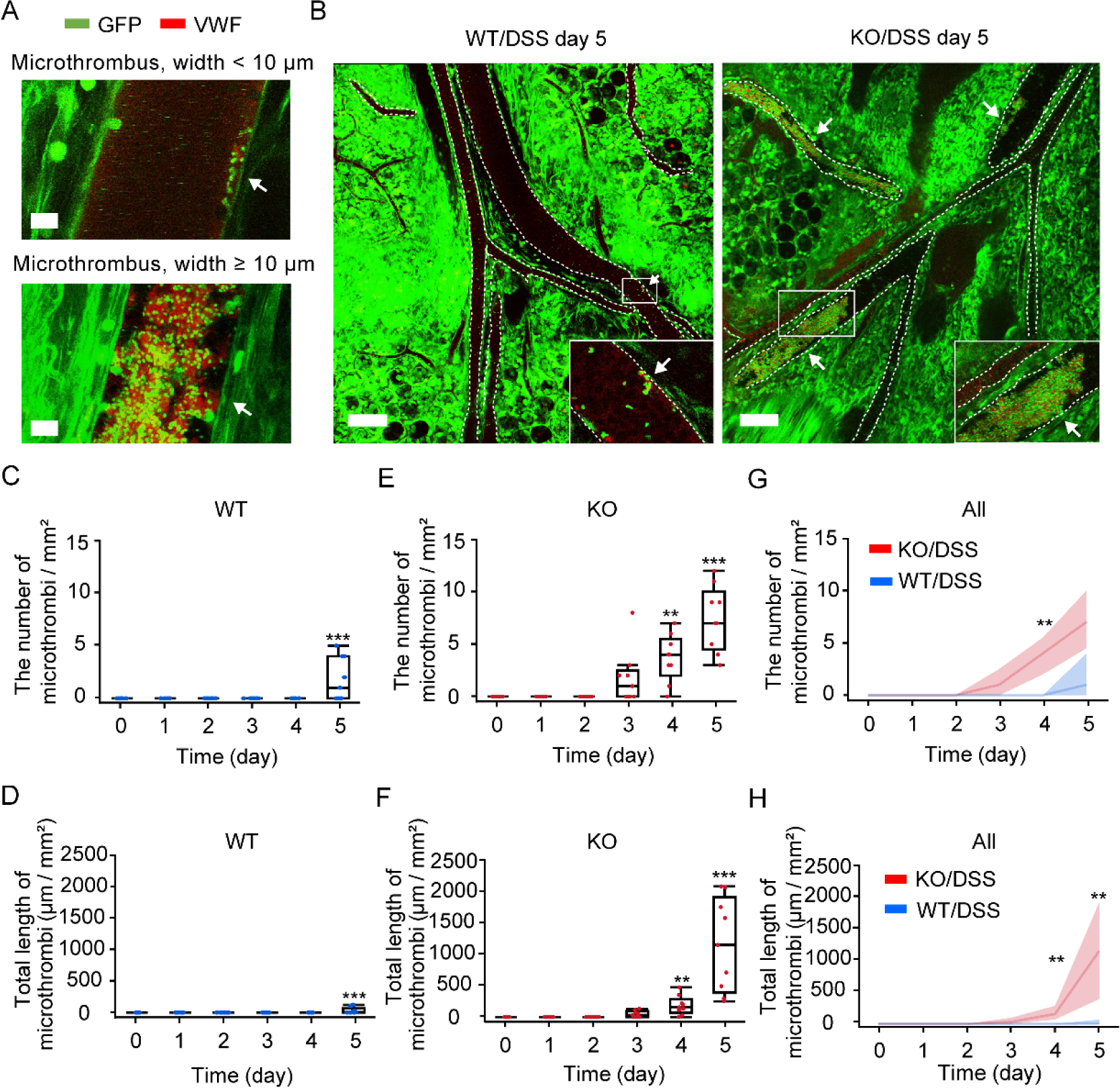
Microvascular thrombus formation in the colonic mucosal and submucosal area in DSS-induced colitis. Mice (age: 7–8 weeks, male, n=9 per group) were administered 5% DSS for 5 days. Thirty minutes before intravital observation, Alexa Fluor 568-labeled anti-VWF antibody was injected intravenously. (A) Representative images of microthrombus categorized by thrombus width. Green, GFP; red, anti-VWF antibody; scale bar: 10 µm (B) Representative images of microthrombi within mucosal and submucosal vessels of WT and ADAMTS13-KO mice 5 days after DSS administration (each inset shows the magnified image of the boxed region; white arrows, microthrombi with VWF accumulation; scale bar: 100 µm). (C, E, G) Changes over time in the number of microthrombi in WT mice (blue) (C), ADAMTS13-KO mice (red) (E), and both genotypes of mice (G) 0–5 days after DSS administration in a large observation area (1 × 1 mm field). (D, F, H) Changes in the total length of microthrombi in WT mice (blue) (D), ADAMTS13-KO mice (red) (F), and both genotypes of mice (H) at 0–5 days after DSS administration in a large observation area. Data (C–F) are shown as the median (center line), 25th and 75th percentiles (box bounds), and whiskers (maximum and minimum data points). Statistically significant changes were assessed using Steel’s multiple comparison test to compare values recorded on day 0 with those on the other days. Data (G, H) are presented as a median and interquartile range, with statistical significance assessed using the Mann–Whitney U test with Bonferroni correction. **p<0.01; ***p<0.001.

To compare microthrombus formation between WT and ADAMTS13-KO mice, we analyzed 1 × 1 mm image field for quantified microthrombi number and length from days 0 to 5 of DSS administration. WT mice showed an increase in both parameters on day 5, though to a limited extent (Figure 4C and 4D). In contrast, ADAMTS13-KO mice demonstrated earlier and more extensive microthrombus formation than WT mice with a significant increase from day 4 (Figure 4E-H). A vessel diameter analysis showed that WT mice primarily exhibited small-vessel microthrombi (10–50 µm; Figure S7A and S7C), whereas ADAMTS13-KO mice exhibited widespread microthrombi across vessels of all diameters (Figure S7). These results suggest that ADAMTS13-KO mice experience limited cleavage of endothelial cell-released UL-VWF multimers, triggering increased platelet aggregation and accelerated microthrombus formation in the mucosal and submucosal vasculature. Moreover, leukocyte recruitment was notably enhanced in ADAMTS13-KO mice, particularly in small vessels, as demonstrated by increased rolling counts (Figure S6A and S6C), suggesting a potential link to diminished mucus production from the luminal side.

### Microthrombi in mucosal regions disrupt microcirculation and exacerbate colitis

To explore the impact of microthrombi on colitis progression, we analyzed their distribution within mucosal and submucosal vessels in ADAMTS13-KO mice. We first categorize regional specificity based on their anatomical locations into three regions: the lower part of the crypt (Area I), intermediate regions (Area II), and submucosa (Area III) (Figure 5A; Figure S8A). The submucosal vessels, primarily consisting of arteries and veins, are larger, with diameters exceeding 40 µm. Vessel diameters gradually decrease toward the lower part of the crypt (Figure 5B).

**Figure 5.**
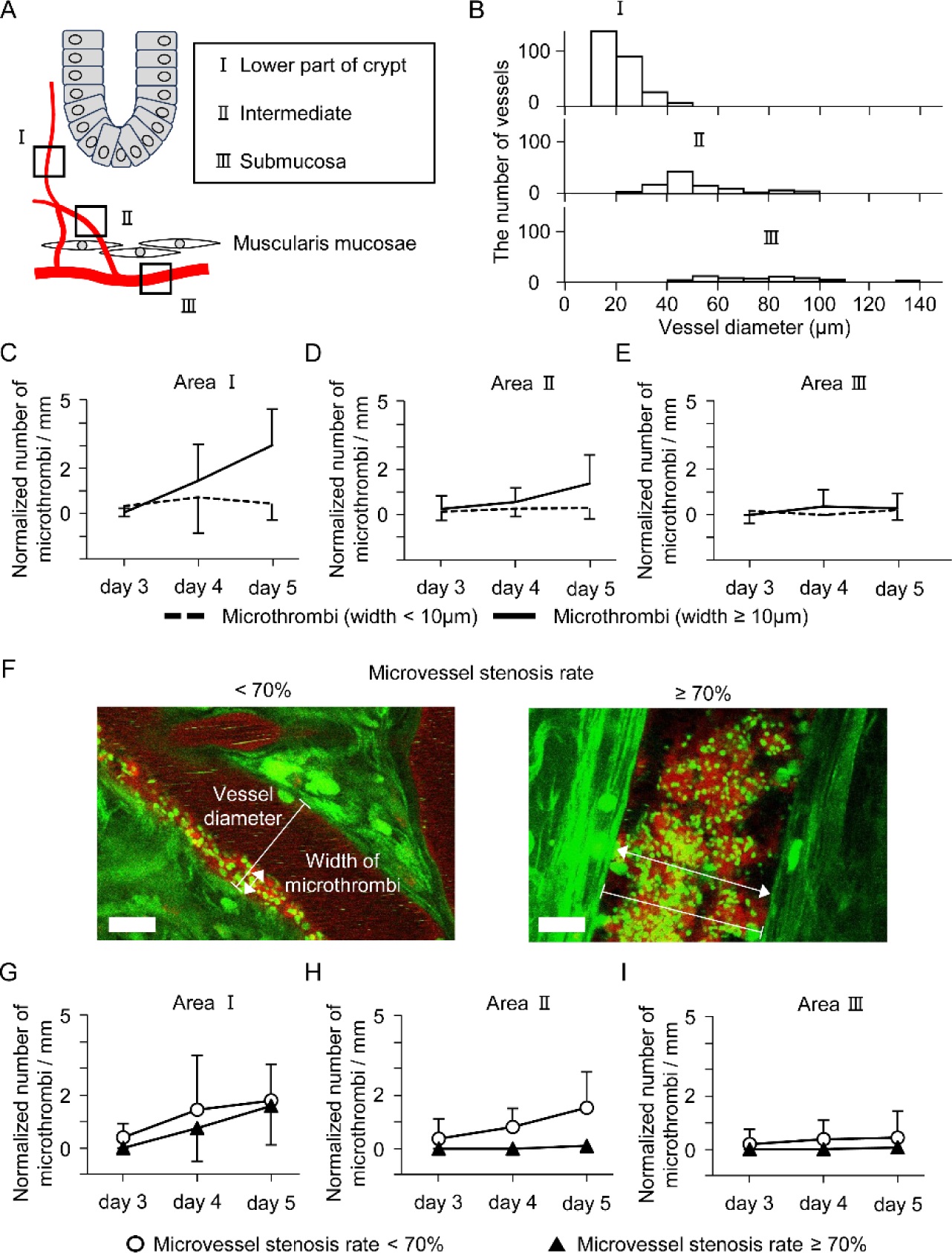
Spatiotemporal analysis of microthrombi in the mucosal and submucosal vessels. ADAMTS13-KO Mice (age: 7–8 weeks, male, n=9) were administered 5% DSS for 5 days, and all observed microthrombi were analyzed. (A) Microvascular anatomical classification of colonic crypts: a. the lower part of the crypt (Area I), b. intermediate area (Area II), c. submucosa (Area III). (B) Distribution of vessel diameters according to the anatomical classification of colonic epithelial. (C-E) Comparison of temporal changes in the normalized number of microthrombi at the 10 µm width boundary in vessels from Area I (C), Area II (D), and Area III (E). The number of microthrombi per millimeter was calculated by normalizing to the length of the blood vessel. Microthrombi with a width of ≥ 10 µm at Area I (C, black line) showed a significant increase over time (p<0.001). Microthrombi with the width < 10µm: dotted black line; microthrombi with the width ≥ 10µm: black line. (F) Calculation method for microvascular stenosis rate. Green, GFP; red, anti-VWF antibody scale bar: 10 µm. (G-I) Comparison of temporal changes in the normalized number of microthrombi at the 70% stenosis rate boundary in the vessels at Area I (G), Area II (H), and Area III (I). Microthrombi with stenosis rate ≥ 70% at Area I (G, black triangle) showed a significant increase over time (p<0.01). Stenosis rates less than 70%, white circle; 70% or more, black triangle. Data (C-E, G-I; mean ± SD) were analyzed by Dunnett’s multiple comparison test to compare values recorded on day 3 with those on the other days.

Microthrombi (< 10 µm width) were detected in vessels across all areas by day 3, with the highest prevalence in Area I, where their presence persisted over time (Figure 5C through 5E). Larger microthrombi (≥ 10 µm width) first appeared in Area I on day 3 and increased markedly by day 5 (Figure 5C), with a moderate presence in Area II (Figure 5D). In contrast, Area III showed minimal microthrombus formation, which remained low throughout the observation period (Figure 5E). These area-specific differences were consistent with vessel diameter classifications shown in Figure S8.

Microthrombi formed in the lower part of the crypt contributes to impaired local blood flow, resulting in mucosal microcirculatory disturbances and potential ischemia. Thrombi that occupy more than 70% of the vessel diameter can significantly restrict or occlude blood flow, as predicted by the Hagen–Poiseuille equation and demonstrated in the previous report.^35^ Stenosis rates were assessed by comparing the largest microthrombus width to the vessel diameter across Areas I–III (Figure 5F). Microthrombi with a stenosis rate ≥70% were observed in Area I beginning on day 4, with a significant increase by day 5 (Figure 5G). No similar increase was noted in Areas II or III (Figure 5H and 5I). These findings indicate that obstructive microthrombi in the lower part of the crypts exacerbate microcirculatory dysfunction, potentially leading to epithelial ischemia and mucosal erosion (Figure 2D, 2F, and 2G).

### rhADAMTS13 administration reduces microthrombus formation and ameliorates DSS-induced colitis

To assess the therapeutic efficacy of rhADAMTS13 in reducing microthrombus formation, daily intravenous administration of rhADAMTS13 was performed in both genotypes of mice during the early phase of DSS-induced colitis. Treatment with rhADAMTS13 significantly improved disease activity (Figure S9B), reduced mucosal erosion (Figure S9D and S9E), and decreased microthrombus formation on day 5 (Figure 6A and 6B). Detailed regional analysis showed that rhADAMTS13 administration in ADAMTS13-KO mice markedly reduced microthrombus formation in Areas I and II, particularly those with high stenosis rates (Figure 6C). Notably, no microthrombi with a stenosis rate ≥70% were observed in any region following rhADAMTS13 treatment (Figure 6C). These findings indicate that rhADAMTS13 effectively suppresses microthrombus formation, improves colonic microcirculation, and contributes to the resolution of mucosal erosion and overall colitis severity.

**Figure 6.**
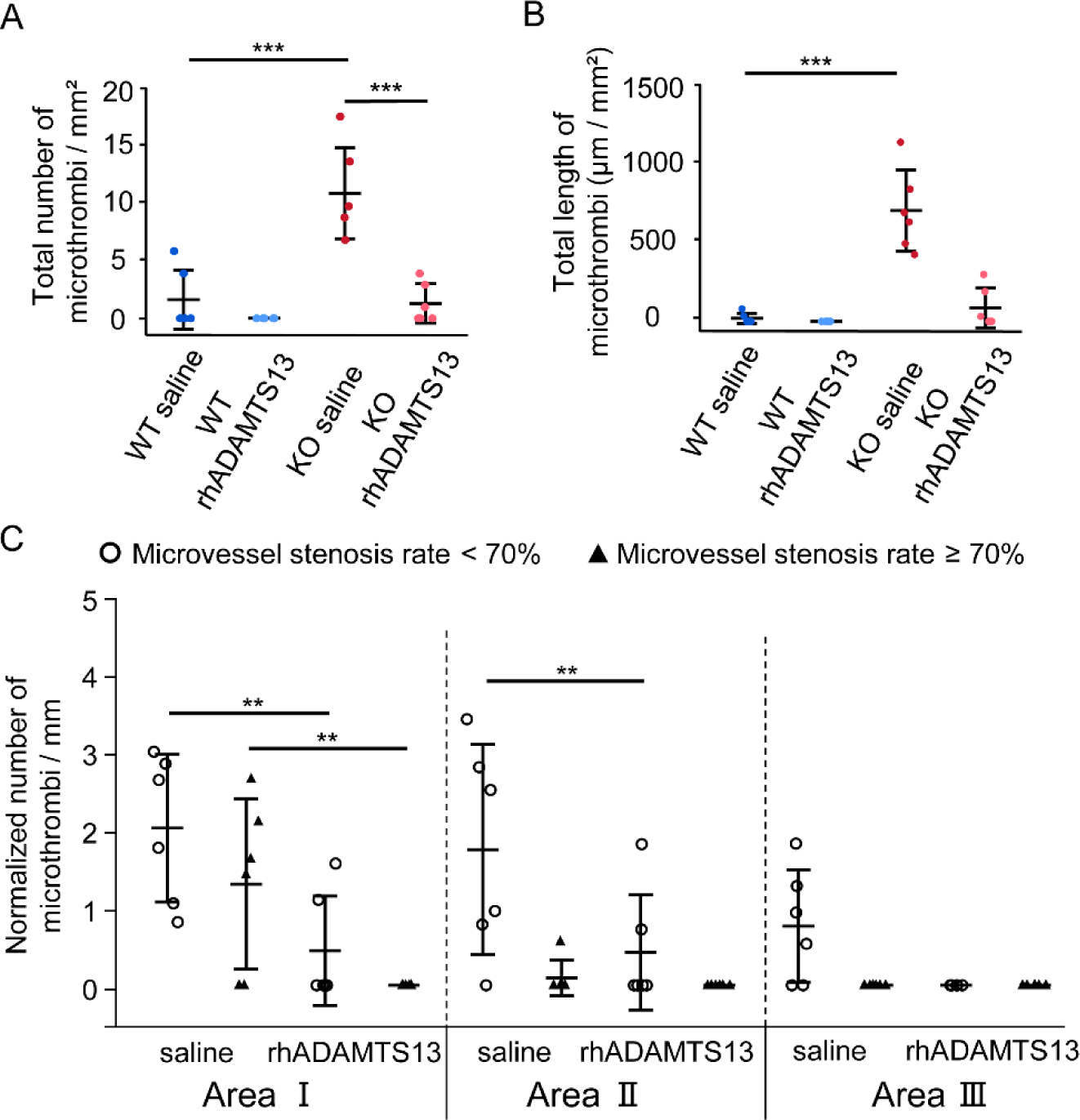
Improvement of mucosal and submucosal vascular microthrombi in DSS-induced colitis after rhADAMTS13 treatment. Mice (age: 7–8 weeks, male, n=6 per group) were administered 5% DSS for 5 days and simultaneously received rhADAMTS13 (WT, light blue dots; KO, light red dots) or saline (WT, blue dots; KO, red dots) injected intravenously on days 1–5. (A) Number of microthrombi on day 5. (B) Total length of microthrombi on day 5. (C) The normalized number of microthrombi on day 5 according to microvascular anatomical classification. Data (A-C; mean ± SD) were analyzed by the Student’s t-test with Bonferroni correction. **p<0.01; ***p<0.001.

## Discussion

In this study, we used advanced intravital imaging techniques with multiphoton excitation microscopy to directly visualize events occurring within the mucosal layer during colitis. We demonstrated for the first time that the early stages of colitis, beginning with mucus layer dysfunction and microthrombus formation in the mucosal vessels, play a critical role in exacerbating disease severity. The mucus layer loses its protective function after DSS administration, with this loss progressing faster in ADAMTS13-KO mice than in WT mice. Concurrently, microthrombi due to ADAMTS13 deficiency enlarge over time, leading to stenosis, particularly in the vessels of the lower part of colonic crypts. This narrowing of the vessels causes a microcirculatory dysfunction, exacerbating colitis. RhADAMTS13 effectively suppresses microthrombus formation, thereby improving the severity of colitis. These findings suggest that a deficiency in ADAMTS13 leads to microthrombus formation in the mucosal vessels, contributing to the refractory nature of UC.

Minimally invasive direct visualization successfully revealed subtle changes in the early stages of colitis pathogenesis. Previous studies of intravital intestinal imaging involved intestinal incisions and other surgical procedures for imaging analysis preparation.^36,37^ Although these invasive procedures effectively mitigate intestinal peristaltic activity, which interferes with imaging analysis, they may also introduce secondary or artificial effects in the tissues and microcirculation. Our approach, using only agarose fixation without intestinal incision, enabled rigorous observation of the functional mucus layer and intestinal microcirculation, allowing for the first-time evidence of microthrombi’s involvement in early-stage UC pathogenesis.

Detailed observation of the functional mucus layer by TRITC-labeled dextran intrusion from the lumen to the epithelium enabled the early detection of alterations following the DSS administration to induce experimental colitis. The mucus layer undergoes constant renewal through mucin secretion from the goblet cells. The estimated mucus layer turnover duration in live murine distal colonic tissue is 1–2 h.^38^ Thus, in vivo assessment of the dynamic changes within the mucus layer, characterized by pronounced fluctuations, is essential to elucidate the pathogenic characteristics of early-stage UC. However, previous studies have evaluated the mucus layer using fixed tissues with Carnoy’s fixative solution^39^ or ex vivo methods,^40^ which makes it challenging to analyze the fluctuating functional structure of mucus layers. Here, by quantifying the distribution of fluorescent dextran in colonic crypts, we introduced a novel approach for real-time functional assessment of the colonic mucus layer that enables the identification of the onset of mucus layer dysfunction during colitis progression.

In UC, disruption of the mucus layer triggers a strong inflammatory response within the intestinal epithelium.^4^ We found that vessels in the lower part of the crypt were more susceptible to thrombus formation than the area under the muscularis mucosae, i.e., the submucosal area. A previous study showed that the aggregation of several aligned platelets within vessels, in response to endothelial activation via topical application of Ca^2+^, resulted in progressive thrombus formation and eventual vascular occlusion.^41–43^ This phenomenon was particularly evident in ADAMTS13-KO mice. In our study, damaged colonic crypts in a colitis pathology model of ADAMTS13-KO mice affected the mucosal vascular endothelium, leading to aligned platelets, followed by the growth of microthrombi in the mucosal vessels, resulting in severe stenosis exceeding 70%. Severe microthrombi-dependent stenosis could induce microcirculatory dysfunction. Previous study has shown that ischemia and ischemia-reperfusion injury lead to mucosal damage.^44^ The absence of microthrombi causing severe stenosis of over 70%, along with the significant improvement in mucosal erosion following rhADAMTS13 administration, strongly supported this evidence. Hence, the ADAMTS13-VWF system plays a significant role in the early stages of pathogenesis and contributes to its exacerbation, depending on the severity of the inflammatory disease.

Recent findings highlight the crucial role of VWF in inflammation, where it recruits leukocytes via direct or platelet-dependent interactions.^45^ In vitro studies show VWF binding to leukocyte P-selectin glycoprotein ligand-1 and β2-integrins,^46^ mimicking interactions with P-selectin and intercellular adhesion molecules for rolling and stable adhesion.^47^ UL-VWF-binding platelets indirectly support leukocyte recruitment by expressing p-selectin on their activated surfaces under high shear stress.^48^ Therefore, in addition to microcirculatory dysfunction, we suggest that VWF-leukocyte interactions during mucosal microvascular thrombus formation promote inflammatory cell recruitment to the colonic mucosa, exacerbating colitis. This mechanism of microvascular thrombosis-induced organ damage is implicated in conditions like deep vein thrombosis,^49^ stroke,^50^ and cardiovascular diseases.^51^

In conclusion, this study provides compelling evidence that the microthrombi formed at the colonic mucosal vessels by insufficient ADAMTS13 activity contributes to exacerbating the early stages of UC pathogenesis. Resolving circulatory dysfunction caused by microthrombus formation is crucial for overcoming the refractory condition of UC.

## Non-standard Abbreviations and Acronyms

UC: ulcerative colitis
VWF: von Willebrand factor
ADAMTS13: a disintegrin-like and metalloproteinase with thrombospondin type 1 motif 1
UL: ultra-large
rhADAMTS13: recombinant human ADAMTS13
GFP: green fluorescent protein
DSS: dextran sulfate sodium DSS
PAS: periodic acid Schiff
TRITC: tetramethylrhodamine
ROI: regions of interest

## Acknowledgments

Fluorescence imaging experiments were performed at the Advanced Research Facilities and Services of the Hamamatsu University School of Medicine.

## Sources of Funding

This work was supported by the JSPS KAKENHI [grant numbers JP22K16486 (M.S.) and JP22K08153 (Y.S.)], a Grant-in-Aid for Scientific Research (B) [grant number JP21H03352 (N.H.) and JP24K02857 (N.H.)], JST PRESTO [grant number JPMJPR19G9 (N.H.)], the Takeda Science Foundation (N.H.), and an HUSM Grant-in-Aid (K.T., N.H., Y.S.).

## Disclosures

The authors declared no competing interests.

## Highlights

Intravital imaging with single-cell resolution revealed mucosal structure disruption and area-specific microthrombus formation in ulcerative colitis model.

ADAMTS13 deficiency enhances microvascular obstructive thrombi causing local ischemia and mucosal erosion.

Advanced imaging techniques provid new insights into ulcerative colitis pathophysiology and contribute to resolving the challenges of its refractory condition.

## Notes

### Competing Interest Statement

The authors have declared no competing interest.

### Summary of Updates

Supplemental files (Materials and Methods, 9 Supplemental Figures) have been uploaded.

## References

1. Ungaro R, Mehandru S, Allen PB, Peyrin-Biroulet L, Colombel JF. Ulcerative colitis. Lancet. 2017;389(10080):1756-1770. doi:10.1016/S0140-6736(16)32126-2

2. Van Der Post S, Jabbar KS, Birchenough G, Arike L, Akhtar N, Sjovall H, Johansson MEV, Hansson GC. Structural weakening of the colonic mucus barrier is an early event in ulcerative colitis pathogenesis. Gut. 2019;68(12):2142–2151. doi:10.1136/gutjnl-2018-317571

3. Neurath MF. Cytokines in inflammatory bowel disease. Nat Rev Immunol. 2014;14(5):329–342. doi:10.1038/nri3661

4. Le Berre C, Honap S, Peyrin-Biroulet L. Ulcerative colitis. Lancet. 2023;402(10401):571-584. doi:10.1016/S0140-6736(23)00966-2

5. Sandborn WJ, Su C, Sands BE, D’Haens GR, Vermeire S, Schreiber S, Danese S, Feagan BG, Reinisch W, Niezychowski W, Friedman G, Lawendy N, Yu D, Woodworth D, Mukherjee A, Zhang H, Healey P, Panés J. Tofacitinib as induction and maintenance therapy for ulcerative colitis. New Engl J Med. 2017;376(18):1723–1736. doi:10.1056/nejmoa1606910

6. Sands BE, Peyrin-Biroulet L, Loftus E V., Danese S, Colombel JF, Törüner M, Jonaitis L, Abhyankar B, Chen J, Rogers R, Lirio RA, Bornstein JD, Schreiber S. Vedolizumab versus adalimumab for moderate-to-severe ulcerative colitis. New Engl J Med. 2019;381(13):1215–1226. doi:10.1056/nejmoa1905725

7. Deban L, Correale C, Vetrano S, Malesci A, Danese S. Multiple Pathogenic Roles of microvasculature in inflammatory bowel disease: A jack of all trades. Am J Pathol. 2008;172(6):1457–1466. doi:10.2353/ajpath.2008.070593

8. Stevens TRJ, James JP, Simmonds NJ, McCarthy DA, Laurenson IF, Maddison PJ, Rampton DS. Circulating von Willebrand factor in inflammatory bowel disease. Gut. 1992;33(4):502–506. doi:10.1136/gut.33.4.502

9. Crawley JTB, de Groot R, Xiang Y, Luken BM, Lane DA. Unraveling the scissile bond: how ADAMTS13 recognizes and cleaves von Willebrand factor. Blood. 2011;118(12):3212–3221. doi:10.1182/blood-2011-02-306597

10. Bernardo A, Ball C, Nolasco L, Moake JF, Dong JF. Effects of inflammatory cytokines on the release and cleavage of the endothelial cell-derived ultralarge von Willebrand-factor multimers under flow. Blood. 2004;104(1):100–106. doi:10.1182/blood-2004-01-0107

11. Cibor D, Owczarek D, Butenas S, Salapa K, Mach T, Undas A. Levels and activities of von Willebrand factor and metalloproteinase with thrombospondin type-1 motif, number 13 in inflammatory bowel diseases. World J Gastroenterol. 2017;23(26):4796–4805. doi:10.3748/wjg.v23.i26.4796

12. Zitomersky N, Demers M, Martinod K, Gallant M, Cifuni S, Biswas A, Snapper S, Wagner D. ADAMTS13 deficiency worsens colitis and exogenous ADAMTS13 administration decreases colitis severity in mice. TH Open. 2017;01(01):e11–e23. doi:10.1055/s-0037-1603927

13. Dhillon AP, Anthony A, Sim R, Wakefield AJ, Sankey EA, Hudson M, Allison MC, Pounder RE. Mucosal capillary thrombi in rectal biopsies. Histopathology. 1992;21(2):127–133. doi:10.1111/j.1365-2559.1992.tb00360.x

14. He G, Ouyang Q, Chen D, Li F, Zhou J. The microvacular thrombi of colonic tissue in ulcerative colitis. Dig Dis Sci. 2007;52(9):2236–2240. doi:10.1007/s10620-006-9158-5

15. Suzuki Y, Mogami H, Ihara H, Urano T. Unique secretory dynamics of tissue plasminogen activator and its modulation by plasminogen activator inhibitor-1 in vascular endothelial cells. Blood. 2009;113:470–478. doi:10.1182/blood-2008-03

16. Suzuki Y, Yasui H, Brzoska T, Mogami H, Urano T. Surface-retained tPA is essential for effective fibrinolysis on vascular endothelial cells. Blood. 2011;118(11):3182–3185. doi:10.1182/blood-2011-05-353912

17. Brzoska T, Suzuki Y, Sano H, Suzuki S, Tomczyk M, Tanaka H, Urano T. Imaging analyses of coagulation-dependent initiation of fibrinolysis on activated platelets and its modification by thrombin-activatable fibrinolysis inhibitor. Thromb Haemost. 2017;117(04):682–690. doi:10.1160/TH16-09-0722

18. Falati S, Gross P, Merrill-Skoloff G, Furie BC, Furie B. Real-time in vivo imaging of platelets, tissue factor and fibrin during arterial thrombus formation in the mouse. Nat Med. 2002;8(10):1175–1180. doi:10.1038/nm782

19. Brzoska T, Tanaka-Murakami A, Suzuki Y, Sano H, Kanayama N, Urano T. Endogenously generated plasmin at the vascular wall injury site amplifies lysine binding site-dependent plasminogen accumulation in microthrombi. PLoS One. 2015;10(3):e0122196. doi:10.1371/journal.pone.0122196

20. Hayashi T, Mogami H, Murakami Y, Nakamura T, Kanayama N, Konno H, Urano T. Real-time analysis of platelet aggregation and procoagulant activity during thrombus formation in vivo. Pflugers Arch. 2008;456(6):1239–1251. doi:10.1007/s00424-008-0466-9

21. Rybaltowski M, Suzuki Y, Mogami H, Chlebinska I, Brzoska T, Tanaka A, Banno F, Miyata T, Urano T. In vivo imaging analysis of the interaction between unusually large von Willebrand factor multimers and platelets on the surface of vascular wall. Pflugers Arch. 2011;461(6):623–633. doi:10.1007/s00424-011-0958-x

22. Honkura N, Richards M, Laviña B, Sáinz-Jaspeado M, Betsholtz C, Claesson-Welsh L. Intravital imaging-based analysis tools for vessel identification and assessment of concurrent dynamic vascular events. Nat Commun. 2018;9(1):2746. doi:10.1038/s41467-018-04929-8

23. Okabe M, Ikawa M, Kominami K, Nakanishi T, Nishimune Y. “Green mice” as a source of ubiquitous green cells. FEBS Lett. 1997;407(3):313–319. doi:10.1016/S0014-5793(97)00313-X

24. Banno F, Kokame K, Okuda T, Honda S, Miyata S, Kato H, Tomiyama Y, Miyata T. Complete deficiency in ADAMTS13 is prothrombotic, but it alone is not sufficient to cause thrombotic thrombocytopenic purpura. Blood. 2006;107(8):3161–3166. doi:10.1182/blood-2005-07-2765

25. Kim JJ, Shajib MS, Manocha MM, Khan WI. Investigating intestinal inflammation in DSS-induced model of IBD. J Vis Exp. 2012;(60):1–6. doi:10.3791/3678

26. Schroeder KW. Coated oral 5-aminosalicylic acid therapy for mildly to moderately active ulcerative colitis. A randomized study. N Engl J Med. 1987;317(26):1625–1629.

27. Sands BE. Biomarkers of inflammation in inflammatory bowel disease. Gastroenterology. 2015;149(5):1275–1285.e2. doi:10.1053/j.gastro.2015.07.003

28. Haberman Y, Karns R, Dexheimer PJ, Schirmer M, Somekh J, Jurickova I, Braun T, Novak E, Bauman L, Collins MH, Mo A, Rosen MJ, Bonkowski E, Gotman N, Marquis A, Nistel M, Rufo PA, Baker SS, Sauer CG, Markowitz J, Pfefferkorn MD, Rosh JR, Boyle BM, Mack DR, Baldassano RN, Shah S, Leleiko NS, Heyman MB, Grifiths AM, Patel AS, Noe JD, Aronow BJ, Kugathasan S, Walters TD, Gibson G, Thomas SD, Mollen K, Shen-Orr S, Huttenhower C, Xavier RJ, Hyams JS, Denson LA. Ulcerative colitis mucosal transcriptomes reveal mitochondriopathy and personalized mechanisms underlying disease severity and treatment response. Nat Commun. 2019;10(1):38. doi:10.1038/s41467-018-07841-3

29. Fenton CG, Taman H, Florholmen J, Sørbye SW, Paulssen RH. Transcriptional signatures that define ulcerative colitis in remission. Inflamm Bowel Dis. 2021;27(1):94–105. doi:10.1093/ibd/izaa075

30. Okayasu I, Hatakeyama S, Yamada M, Ohkusa T, Inagaki Y, Nakaya R. A novel method in the induction of reliable experimental acute and chronic ulcerative colitis in mice. Gastroenterology. 1990;98(3):694–702. doi: 10.1016/0016-5085(90)90290-h.

31. Wirtz S, Popp V, Kindermann M, Gerlach K, Weigmann B, Fichtner-Feigl S, Neurath MF. Chemically induced mouse models of acute and chronic intestinal inflammation. Nat Protoc. 2017;12(7):1295–1309. doi:10.1038/nprot.2017.044

32. Johansson ME V., Phillipson M, Petersson J, Velcich A, Holm L, Hansson GC. The inner of the two Muc2 mucin-dependent mucus layers in colon is devoid of bacteria. Proc Natl Acad Sci U S A. 2008;105(39):15064–15069. doi:10.1073/pnas.0803124105

33. Grootjans J, Hundscheid IHR, Lenaerts K, Boonen B, Renes IB, Verheyen FK, Dejong CH, von Meyenfeldt MF, Beets GL, Buurman WA. Ischaemia-induced mucus barrier loss and bacterial penetration are rapidly counteracted by increased goblet cell secretory activity in human and rat colon. Gut. 2013;62(2):250–258. doi:10.1136/gutjnl-2011-301956

34. Zhang D, Frenette PS. Cross talk between neutrophils and the microbiota. Blood. 2019;133(20):2168–2177. doi:10.1182/blood-2018-11-844555

35. Sato M, Ohshima N. Hemodynamics at stenoses formed by growing platelet thrombi in mesenteric microvasculature of rat. Microvasc Res. 1986;31(1):66–76. doi:10.1016/0026-2862(86)90007-5

36. Gustafsson JK, Davis JE, Rappai T, McDonald KG, Kulkarni DH, Knoop KA, Hogan SP, Fitzpatrick JA, Lencer WI, Newberry RD. Intestinal goblet cells sample and deliver lumenal antigens by regulated endocytic uptake and transcytosis. Elife. 2021;10;e67292. doi:10.7554/eLife.67292

37. Sullivan DP, Bui T, Muller WA, Butin-Israeli V, Sumagin R. In vivo imaging reveals unique neutrophil transendothelial migration patterns in inflamed intestines. Mucosal Immunol. 2018;11(6):1571–1581. doi:10.1038/s41385-018-0069-5

38. Johansson ME V. Fast renewal of the distal colonic mucus layers by the surface goblet cells as measured by in vivo labeling of mucin glycoproteins. PLoS One. 2012;7(7):e41009. doi:10.1371/journal.pone.0041009

39. Naama M, Telpaz S, Awad A, Ben-Simon S, Harshuk-Shabso S, Modilevsky S, Rubin E, Sawaed J, Zelik L, Zigdon M, Asulin N, Turjeman S, Werbner M, Wongkuna S, Feeney R, Schroeder BO, Nyska A, Nuriel-Ohayon M, Bel S. Autophagy controls mucus secretion from intestinal goblet cells by alleviating ER stress. Cell Host Microbe. 2023;31(3):433–446.e4. doi:10.1016/j.chom.2023.01.006

40. Gustafsson JK, Ermund A, Johansson ME V, Schütte A, Hansson GC, Sjövall H. An ex vivo method for studying mucus formation, properties, and thickness in human colonic biopsies and mouse small and large intestinal explants. Am J Physiol Gastrointest Liver Physiol. 2012;302:430–438. doi:10.1152/ajpgi.00405.2011.

41. Chauhan AK, Motto DG, Lamb CB, Bergmeier W, Dockal M, Plaimauer B, Scheiflinger F, Ginsburg D, Wagner DD. Systemic antithrombotic effects of ADAMTS13. J Exp Med. 2006;203(3):767–776. doi:10.1084/jem.20051732

42. Adili R, Holinstat M. Formation and resolution of pial microvascular thrombosis in a mouse model of thrombotic thrombocytopenic purpura. Arterioscler Thromb Vasc Biol. 2019;39(9):1817–1830. doi:10.1161/ATVBAHA.119.312848

43. Chauhan AK, Goerge T, Schneider SW, Wagner DD. Formation of platelet strings and microthrombi in the presence of ADAMTS-13 inhibitor does not require P-selectin or β3 integrin. J Thromb Haemost. 2007;5(3):583–589. doi:10.1111/j.1538-7836.2007.02361.x

44. Ikeda H, Suzuki Y, Suzuki M, Koike M, Tamura J, Tong J, Nomura M, Itoh G. Apoptosis is a major mode of cell death caused by ischaemia and ischaemia/reperfusion injury to the rat intestinal epithelium. Gut. 1998;42(4):530–537. doi:10.1136/gut.42.4.530

45. Kawecki C, Lenting PJ, Denis C V. von Willebrand factor and inflammation. J Thromb Haemost. 2017;15(7):1285-1294. doi:10.1111/jth.13696

46. Pendu R, Terraube V, Christophe OD, Gahmberg CG, De Groot PG, Lenting PJ, Denis C V. P-selectin glycoprotein ligand 1 and β2-integrins cooperate in the adhesion of leukocytes to von Willebrand factor. Blood. 2006;108(12):3746–3752. doi:10.1182/blood-2006-03-010322

47. McEver R. Adhesive Interactions of Leukocytes, Platelets, and the Vessel Wall during Hemostasis and Inflammation. Thromb Haemost. 2001;86(09):746–756. doi:10.1055/s-0037-1616128

48. Bernardo A, Ball C, Nolasco L, Choi H, Moake JL, Dong JF. Platelets adhered to endothelial cell-bound ultra-large von Willebrand factor strings support leukocyte tethering and rolling under high shear stress. J Thromb Haemost. 2005;3(3):562–570. doi:10.1111/j.1538-7836.2005.01122.x

49. Pagliari MT, Cairo A, Boscarino M, Mancini I, Pappalardo E, Bucciarelli P, Martinelli I, Rosendaal FR, Peyvandi F. Role of ADAMTS13, VWF and F8 genes in deep vein thrombosis. Miyata T, ed. PLoS One. 2021;16(10):e0258675. doi:10.1371/journal.pone.0258675

50. Xu H, Cao Y, Yang X, Cai P, Kang L, Zhu X, Luo H, Lu L, Wei L, Bai X, Zhu Y, Zhao BQ, Fan W. ADAMTS13 controls vascular remodeling by modifying VWF reactivity during stroke recovery. Blood. 2017;130(1):11–22. doi:10.1182/blood-2016-10-747089

51. De Meyer SF, Savchenko AS, Haas MS, Schatzberg D, Carroll MC, Schiviz A, Dietrich B, Rottensteiner H, Scheiflinger F, Wagner DD. Protective anti-inflammatory effect of ADAMTS13 on myocardial ischemia/reperfusion injury in mice. Blood. 2012;120(26):5217–5223. doi:10.1182/blood-2012-06-439935

