## Supplemental Materials and Methods for "Microthrombi-growth in ADAMTS13 deficiency exacerbated ulcerative colitis via mucosal and endothelial dysfunction"

#### Supplemental Material and Methods

##### Human sample preparations

We used samples from patients with U and healthy volunteers for blood plasma analysis. Blood samples were obtained at Hamamatsu University Hospital between November 2021 and July 2023. For immunohistochemical analysis, formalin-fixed paraffin embedded colon tissues were classified as follows: For the “UC” tissues, the site with the highest degree of inflammation was used. For the “normal colon” tissues, we used the non-cancerous colonic tissues in colorectal cancer specimens as an alternative and examined the most distant area from the tumor. These specimens were derived from patients diagnosed as pathological Stage 0 or I according to the 8th edition of the Union for International Cancer Control-Tumor Node Metastasis classification. Tissue samples were obtained from surgical specimens collected at the Hamamatsu University Hospital between January 2017 and June 2022.

##### Antibodies and reagents

Polyclonal rabbit anti-human von Willebrand factor (VWF) antibody (A0082) was purchased from Dako Cytomation (Glostrup, Denmark) and labelled with Alexa Fluor 568 (Molecular Probes, Eugene, OR, USA) using a protein labeling kit, according to the manufacturer’s instructions. The monoclonal Mouse anti-human VWF antibody (M061601) was purchased from Dako Cytomation (Glostrup, Denmark). Tetramethylrhodamine (TRITC)-dextran (molecular weight: 40,000 Da) was purchased from Sigma-Aldrich (St. Louis, MO). TRITC-dextran (molecular weight: 2,000,000 Da) was purchased from Tdb Labs AB (Uppsala, Sweden). The REAL EnVision Detection System and Peroxidase/DAB Rabbit/Mouse HRP (K500711) were purchased from Dako Cytomation (Glostrup, Denmark). Dextran sulfate sodium (DSS, molecular weight: 36,000–50,000 Da) was purchased from MP Biomedicals (OH, USA.). Agarose (50080) was purchased from Lonza (Basel, Switzerland). The recombinant human ADAMTS13 (4245-ADA-02) was purchased from R&D Systems.

**Definition of UC disease severity**

The disease severity of UC was assessed using the Endoscopic Mayo Score (EMS), where a rating of EMS2 was assigned based on colonoscopic findings, which included marked erythema, absent vascular patterns, friability, and erosions. The detection of spontaneous bleeding and ulceration during colonoscopy resulted in an EMS3 rating. EMS2 was classified as moderate and EMS3 as severe.[30]

**Measurement of plasma ADAMTS13 activity level**

Blood sampling was conducted upon hospitalization or prior to any treatment modifications. The collected blood samples with 3.2% trisodium citrate were centrifuged at 1500 × g for 15 minutes at 4°C. Supernatants were collected as plasma samples and preserved at −80°C. Plasma ADAMTS13 activity was measured using a standard enzyme assay based on monoclonal antibodies generated by cleavage of the GST-VWF73-His fusion protein (SRL, Inc. Tokyo, Japan).

**Immunohistochemical analysis of human tissues**

Sections of formalin-fixed paraffin embedded tissues at 4-μm thickness were deparaffinized and rehydrated, followed by blocking endogenous peroxidase activity using 3% H2O2 for 5 minutes at room temperature. Antigen retrieval was then conducted by incubating the sections in citrate buffer for 20 minutes at 95°C. After cooling to room temperature, the sections were incubated overnight at 4°C with a primary mouse monoclonal antibody against human VWF (M061601, Dako) in antibody diluent (S20202230-2, Dako). The sections were washed in PBS, and the bound antibodies were visualized using HRP-conjugated secondary antibodies and DAB buffer (K500711).

Stained sections were scanned using a NanoZoomer S60 Digital slide scanner (Hamamatsu Photonics, Hamamatsu, Japan). The analysis was conducted on a virtual slide using the HALO digital pathological system (Indica Labs, Tokyo, Japan). The analysis focused on mucosal and submucosal layers across the entire colonic tissue. The HALO algorithm first deconvolves the immunohistochemistry image into hematoxylin and DAB channels, then detects individual cells and their subcellular compartments, that is, nucleus and cell membrane, and scores the cells as high, medium, or low based on the average DAB signal associated with the cell membrane. This analysis evaluated the percentage of all DAB signal intensities, from low to high, across all areas that were analyzed.

**Transcriptome analysis from human database**

GSE117993 and GSE128682 mRNA chips and RNA-seq datasets were downloaded from the NCBI for Biotechnology Information Gene Expression Omnibus repository.[29,30] The collected gene expression data were converted to Log2 value, and VWF messenger ribonucleic acid levels were compared.

**Histological assessment in the DSS-induced colitis mouse**

Sections were stained with Hematoxylin & Eosin and periodic acid-Schiff-positive cells (PAS). Stained sections were scanned using a NanoZoomer S60 Digital slide scanner. The analysis was conducted on a virtual slide using Qupath v. 0.4.3, an open-source software for digital pathology image analysis. Histological assessment was performed using a scoring system for inflammation-associated histological changes in the colon, which was evaluated based on the degree of tissue damage (0, none; 1, isolated focal epithelial damage; 2, mucosal erosions and ulcerations; 3, extensive damage deep into the bowel wall) and inflammatory cell infiltration

(0, infrequent; 1, increased, some neutrophils; 2, submucosal presence of inflammatory cell clusters; 3, transmural cell infiltration) in DSS-induced colitis.[32] Each section of the proximal colon, distal colon, and rectum was individually scored and the average of these scores was calculated. The number and length of the mucosal erosions were quantified across the entire colon, from the proximal colon to the lower rectum. To analyze mucus-producing goblet cells, the number of PAS-positive cells per colonic crypt was counted. For each sample, we determined the number of these cells and the area of the intestinal crypts across 10 colonic crypts and calculated the average number of goblet cells per 100 μm2 of the crypt.

**Intravital imaging system**

A Nikon A1R MP confocal/multiphoton microscope (Nikon Corporation, Tokyo, Japan) with an ultrashort pulse-tunable laser (Ti:$Al_2O_3$ laser, Chameleon; Coherent, Santa Clara, CA, USA) was used for intravital imaging. It was equipped with an LWD Lambda S 40XC WI water-immersion objective lens (Nikon Corporation, Tokyo, Japan). Depending on the image depth, the mean laser power of the sample was between 60 and 80 mW. The image sequences were captured as 12-bit, 512 (or 1024) pixel array images (0.5–1 frame per second). Integrated NIS-Elements software (Nikon Solutions Co., Ltd., Tokyo, Japan) was used to operate the microscope and process raw image data. Two-photon excitation was performed at wavelengths of 960 or 1040 nm, to verify mucus barrier function and mucosal vascular thrombus, respectively, for each experiment. The emitted fluorescence was split using 560-nm and/or 593-nm dichroic mirrors placed in series into green and red channels. The fluorescence was then passed through band-pass emission filters at 525/50 and 629/56 nm and separately collected using GaAsP photomultiplier tubes (Hamamatsu Photonics, Hamamatsu, Japan).

**Image acquisition and analysis**

Mice were anesthetized by intraperitoneal injection of a combination anesthetic agent (medetomidine hydrochloride/midazolam/butorphanol: 0.3/4/5 mg/kg) at a volume of 0.01–0.02 mL/g body weight. To avoid tissue damage, the proximal colon was exposed via a small midline abdominal incision and placed onto a cover glass (thickness 0.13–0.17 μm; Matsunami Glass Ind., Ltd., Kishiwada, Japan) and fixed with 6% agarose (Lonza, Ltd., Basel, Switzerland). The prepared mouse section was placed on a controllable metal heater microscope stage (ThermoPlate, Tokai Hit Co., Ltd., Fujinomiya, Japan).

Images were acquired from the muscle layer up to 30 μm above the end of the colonic crypt in the Z-direction in 3-μm increments. To assess the microthrombi distribution, images with a large field of view of approximately 1 × 1 mm were acquired at similar depths. All data were analyzed using ImageJ or FIJI software (National Institutes of Health and LOCI, University of

Wisconsin, Bethesda, MD, USA).

##### Mucus barrier function

Precisely 15 minutes before imaging, 1 mL of 20 mg/mL TRITC-dextran in saline was administered into the cecum. Diffusion of TRITC-dextran near the observation area was confirmed at the time of imaging (Figure 3A). The region of interest (ROI) size was defined as a circle that had a diameter that was half the short diameter of the colonic crypt (Figure S3A). The average green and red intensities within the ROI were measured for each crypt in the lower and upper one-thirds of the colonic crypts. The extent of TRITC-dextran intrusion was evaluated by determining the ratio of green-to-red intensity. The constant GFP fluorescence intensity detected in the green channel of the detectors originated from almost all cells that drove the CAG promoter in the GFP mice. In particular, the same cell type, even in different individuals, showed very similar GFP expression levels in the image area. Therefore, the ratio of the red intensity was determined based on the green intensity, which has a constant intensity, as the reference. Moreover, it was possible to correct for variability caused by interindividual differences in mice and imaging conditions, such as differences in laser power due to the z-step (Figure 3B). The reference point for TRITC-dextran fluorescence intensity was calculated as the ratio of red-to-green fluorescence in the lower part of the colonic crypts of WT mice before

DSS administration (Day 0). The fluorescence intensity on other days was calculated based on the reference point (Day 0).

**Leukocyte dynamics**

TRITC-dextran in saline was administered via retro-orbital intravenous injection 15 min before imaging. The number of leukocytes rolling and adhesion were manually counted. Arbitrary fields with leukocyte rolling were observed and captured for 60 sec. Leukocyte adhesion was defined by leukocytes that were stationary for >30 sec and were counted per 100-μm vessel length. Analysis was performed using images containing TRITC-dextran (2000 kDa) in blood vessels or images of the green channel (Figure S4A and S4B).

**Thrombus in microvessels**

For visualizing UL-VWF multiple platelets, Alexa Fluor 568-labeled anti-VWF antibody (0.1 mL of approximately 1 mg/mL solution) was intravenously administered. The number and length of the microthrombi were counted in each image (1 mm × 1 mm field). In addition, the number of microthrombi was in detail compared based on the anatomical location of colonic mucosal and submucosal vessels (Figure 5A). The number of thrombi was normalized per millimeter of vessel length. The normalized number of microthrombi, according to vascular anatomical location, was analyzed based on thrombus width (using 10 μm as the threshold, Figure 4A) and the degree of stenosis (calculated by dividing the width of the microthrombus by the diameter of the narrowest part of the vessel, classified as less than 70% or greater than 70%, Figure 5F).

**Statistics**

Statistical analysis was performed using JMP® 16 software (SAS Institute Inc., Cary, NC, USA). The distribution features are presented as mean ± standard deviation or median and interquartile range for variables with skewed distribution or frequency. The principal statistical tests were the Student's t-test and Mann–Whitney U test (for non-Gaussian-distributed data), as appropriate. The normality of the data was assessed using the Shapiro–Wilk test. The Bonferroni correction, Dunnett's multiple comparison test, and Steel's multiple comparison test were used to correct for multiple tests. All the tests were two-tailed. Statistical significance is indicated in each figure legend.

#### Supplemental Figure 1

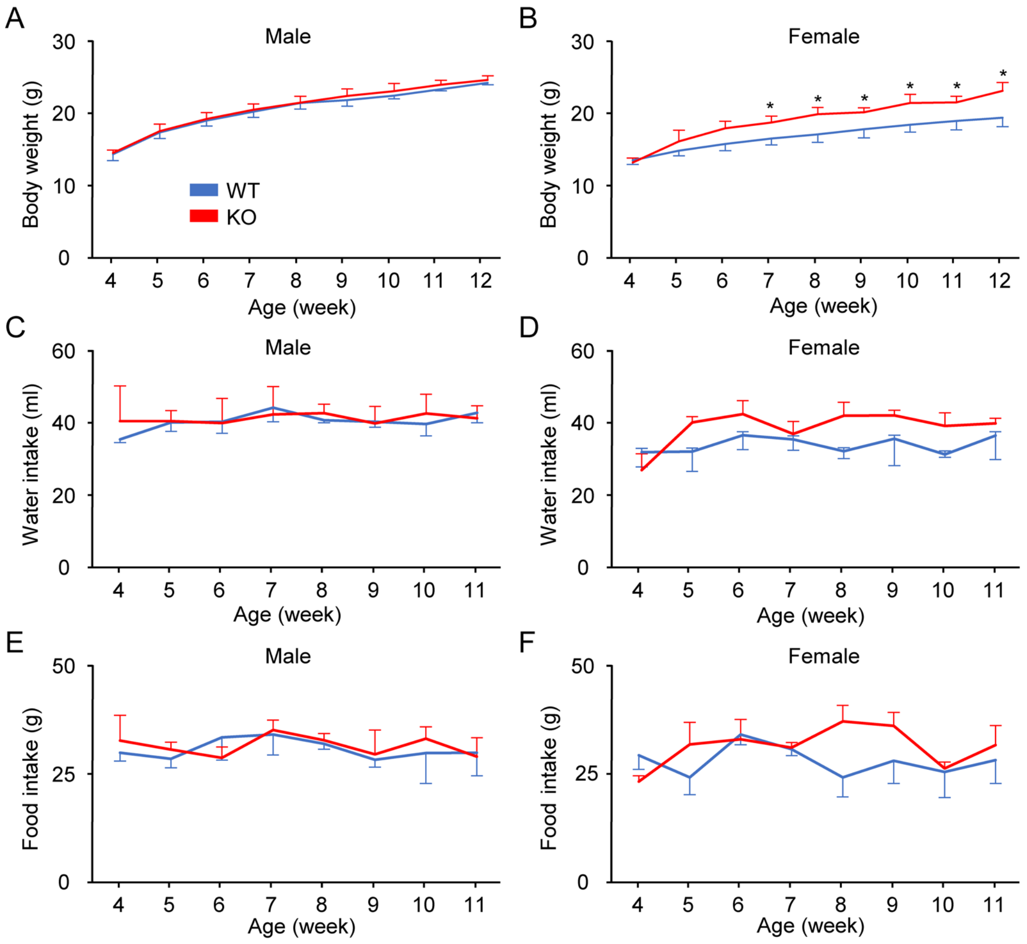


**Supplemental Figure 1**. Development of WT mice and ADAMTS13 knockout mice

The general condition of WT and ADAMTS13-KO mice (n=5 per group) was evaluated from 4 to 12 weeks. Changes in the body weight of male (A) and female (B) mice. Changes in water intake in male (C) and female (D) mice. Changes in food intake in male (E) and female (F) mice. Data was shown as mean and SD. Statistical significance was assessed using the Student's *t*-test with Bonferroni correction. WT, blue lines; KO, red lines. *$p$<0.05.

**Supplemental Material**

#### Supplemental Figure 2

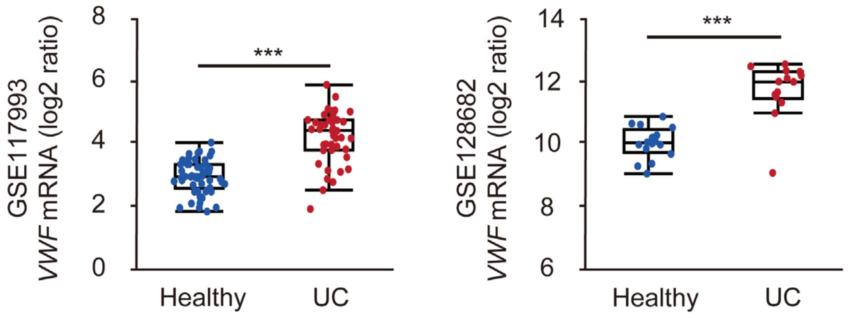


**Supplemental Figure 2. *VWF* mRNA expression in two different Gene Expression Omnibus datasets**

*VWF* mRNA expression levels were obtained from two different Gene Expression Omnibus datasets (GSE117993: Normal: n=55, UC: n=43; GSE 128682: Normal: n=16, UC: n=14). Data are presented as median (center line) with the 25th and 75th percentiles (box bounds) and whiskers (maximum and minimum data points), and were analyzed using the Mann–Whitney *U* test. ***$p$<0.001

#### Supplemental Figure 3

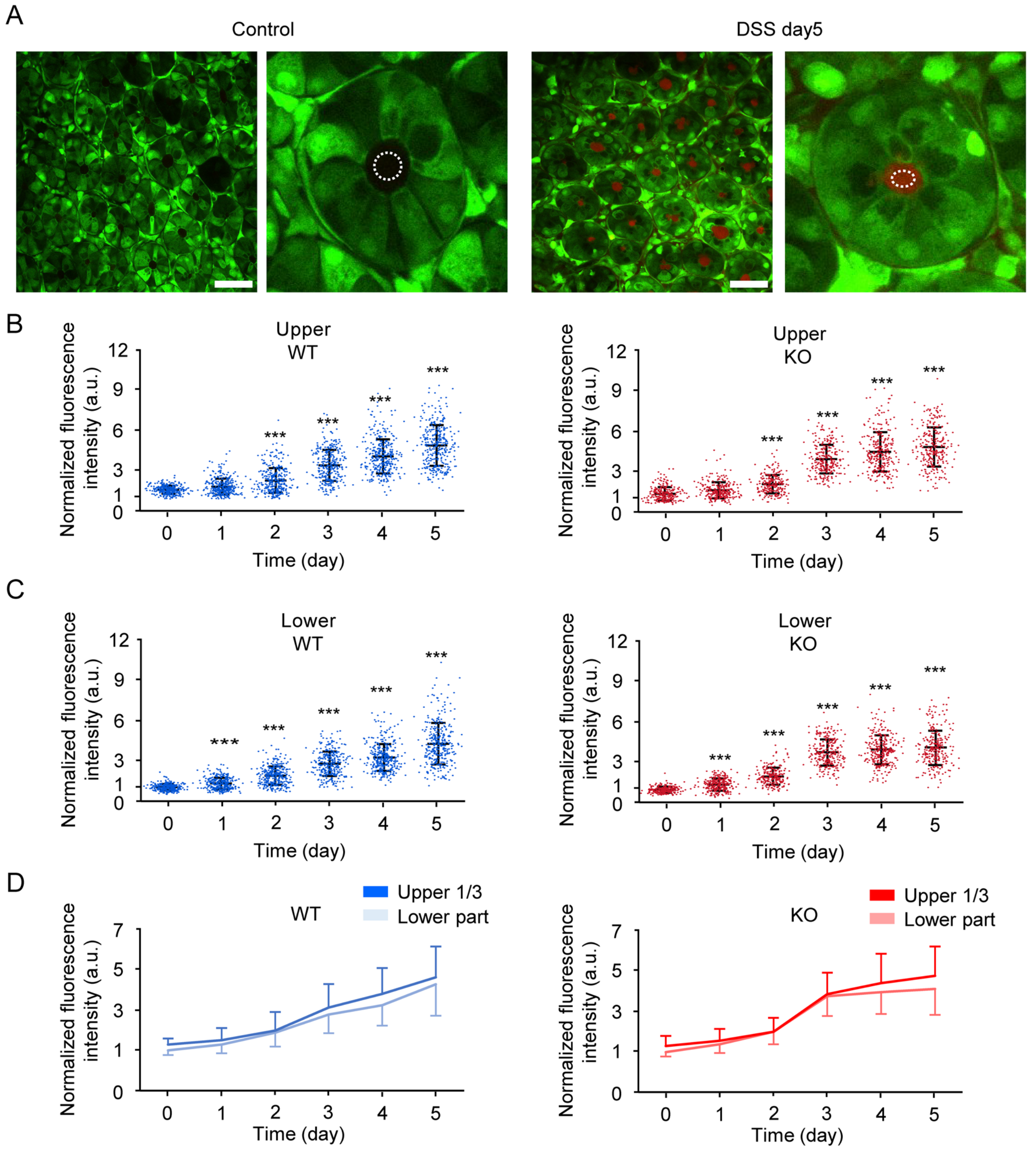


**Supplemental Figure 3. Details of the functional analysis of the mucosal barrier**

Mice (age: 7–8 weeks, male, n=9 per group) were administered with 5% DSS for 5 days. TRITC dextran (40 kDa) was injected into the lumen of the proximal colon, and images were captured 15 min later. (A) The method for quantifying the distribution of TRITC-dextran. The

ROI size (dotted circle) was defined as a circle with a diameter that was half the short diameter of the colonic crypt. Scale bar: 50 μm (B, C) The normalized fluorescence intensity of TRITC-dextran in each ROI in DSS-administered WT mice (blue) and DSS-administered ADAMTS13-KO mice (red) is shown as mean and SD during 0–5 days. (B) Upper one third of the colonic crypts. (C) Lower part of the colonic crypts. (D) Comparison of normalized fluorescence intensity changes according to the anatomical location of the colonic crypts. Light blue lines, lower part of the crypts of WT mice; blue lines, upper one third of the colonic crypts of WT mice; light red lines, lower part of the colonic crypts of ADAMTS13-KO mice; red lines, upper one third of the colonic crypts of ADAMTS13-KO mice. Data (B, C) were assessed by comparing the values recorded on Day 0 with those of the other days using Dunnett's multiple comparison test. Data (D) presented as mean and standard deviation (SD), and analyzed by the Student *t*-test with Bonferroni correction. ***$p<0.001$.

**Supplemental Figure 4**

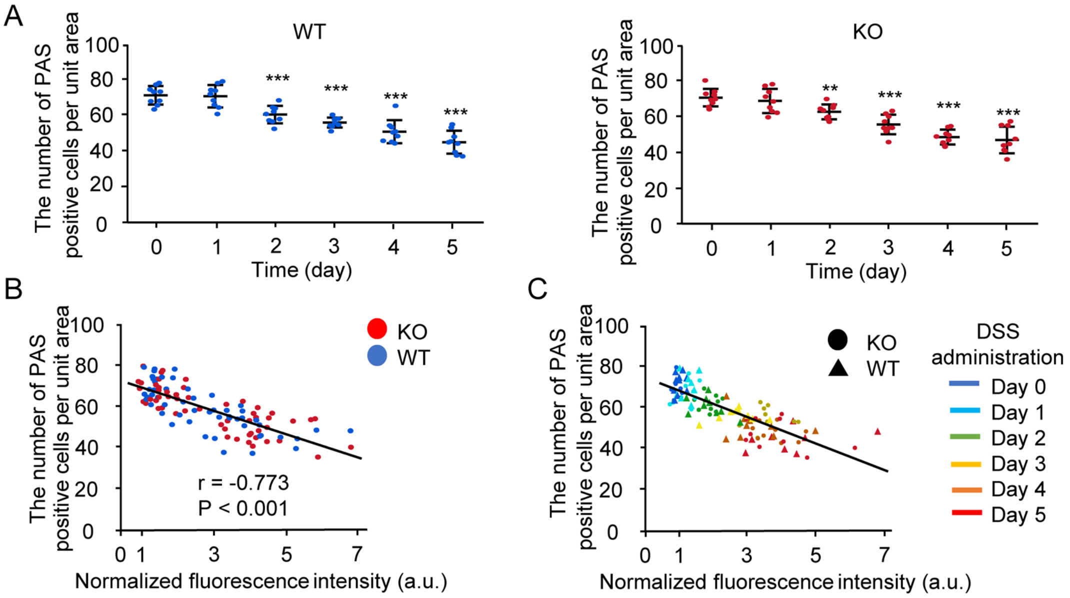


**Supplemental Figure 4. Relationship between the mucus barrier function and goblet cells**

Mice (age: 7–8 weeks, male, n=9 per group) were administered 5% DSS for 5 days. (A) Number of PAS-positive cells per unit area on days 0–5 Blue dots indicate WT mice; red dots indicate ADAMTS13-KO mice. (B) Correlation between the normalized fluorescence intensity in the upper one third of the crypts and the number of goblet cells. Blue dots indicate WT mice; red dots indicate ADAMTS13-KO mice. (C) Correlation between normalized fluorescence intensity in the lower crypts and the number of goblet cells according to the duration of DSS administration. Triangles: WT mice; circles: ADAMTS13-KO mice. Blue, Day 0 of DSS administration; light blue, Day 1; green, Day 2; yellow, Day 3; orange, Day 4; and red, Day 5. Data (A) were represented as mean and SD and analyzed by Dunnett's multiple comparison test. Statistical significance was analyzed using two-sided Spearman's correlation (B, C). $^{**}p<0.01$; $^{***}p<0.001$.

**Supplemental Figure 5**

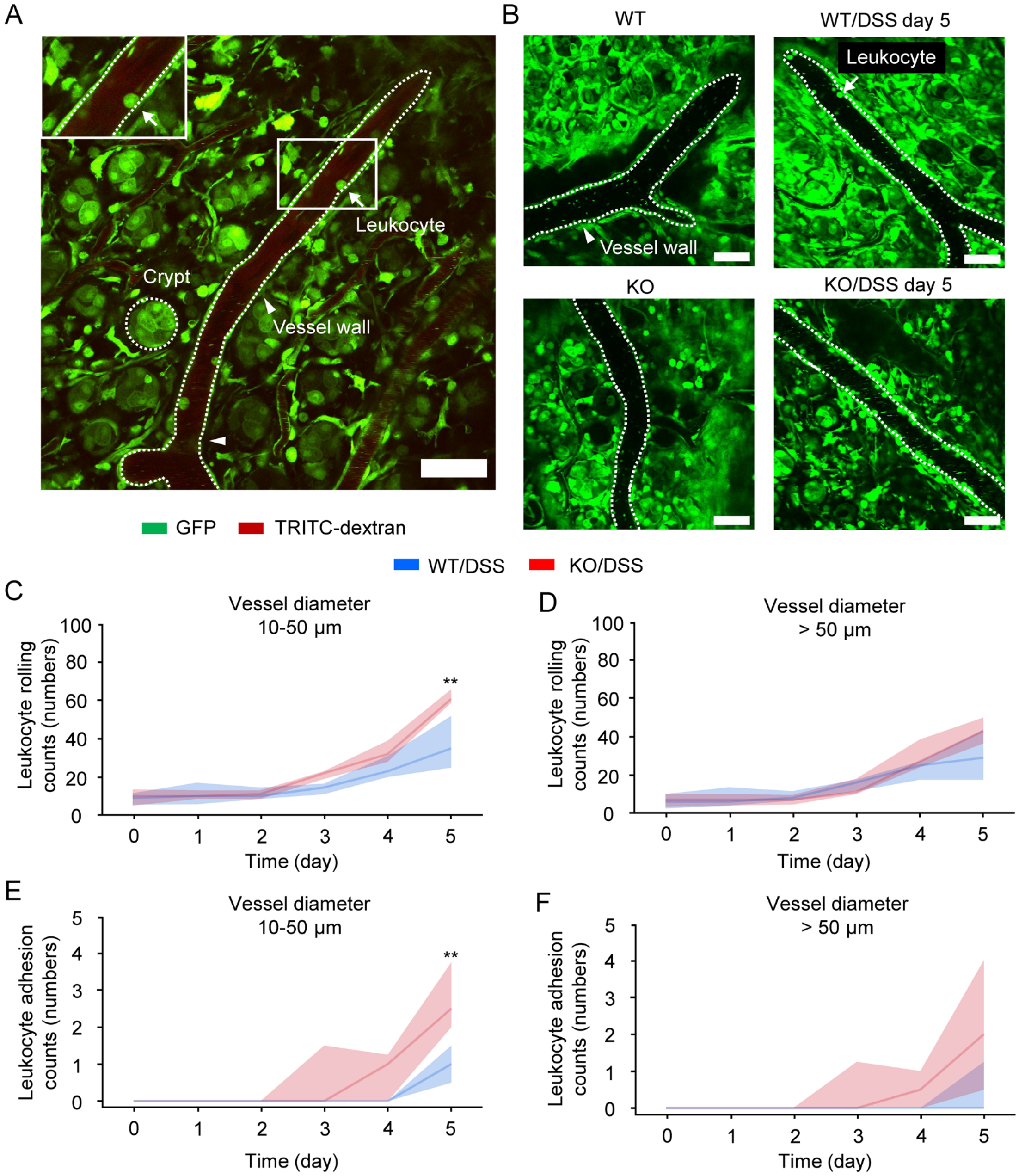


**Supplemental Figure 5. Leukocyte recruitment in DSS-induced colitis**

Mice (age: 7–8 weeks, male, n=9 per group) were administered 5% DSS for 5 days. (A, B) Intravital microscopy-based representative images of mucosal vessels in the proximal colon of WT and ADAMTS13-KO mice 5 days after DSS administration (inset: magnified image of the boxed region; green, GFP; red, TRITC-dextran; white arrows, leukocytes; white arrowheads, vessel walls; and white dotted circles: colonic crypts, scale bar: 50 μm). (C–F) Frequency (median and interquartile range) of leukocyte rolling (C, D) and adhesion (E, F) quantified for 60 seconds in vessels with diameters of 10–50 μm (C, E) and >50 μm (D, F) at 0 to 5 days (blue lines, WT mice; red lines, ADAMTS13-KO mice). Analysis was performed using images containing TRITC-dextran (2000 kDa) in blood vessels or images of the green channel. Statistical significance was assessed using the Mann–Whitney U test with Bonferroni correction (C-F). **$p<0.01$.

#### Supplemental Figure 6

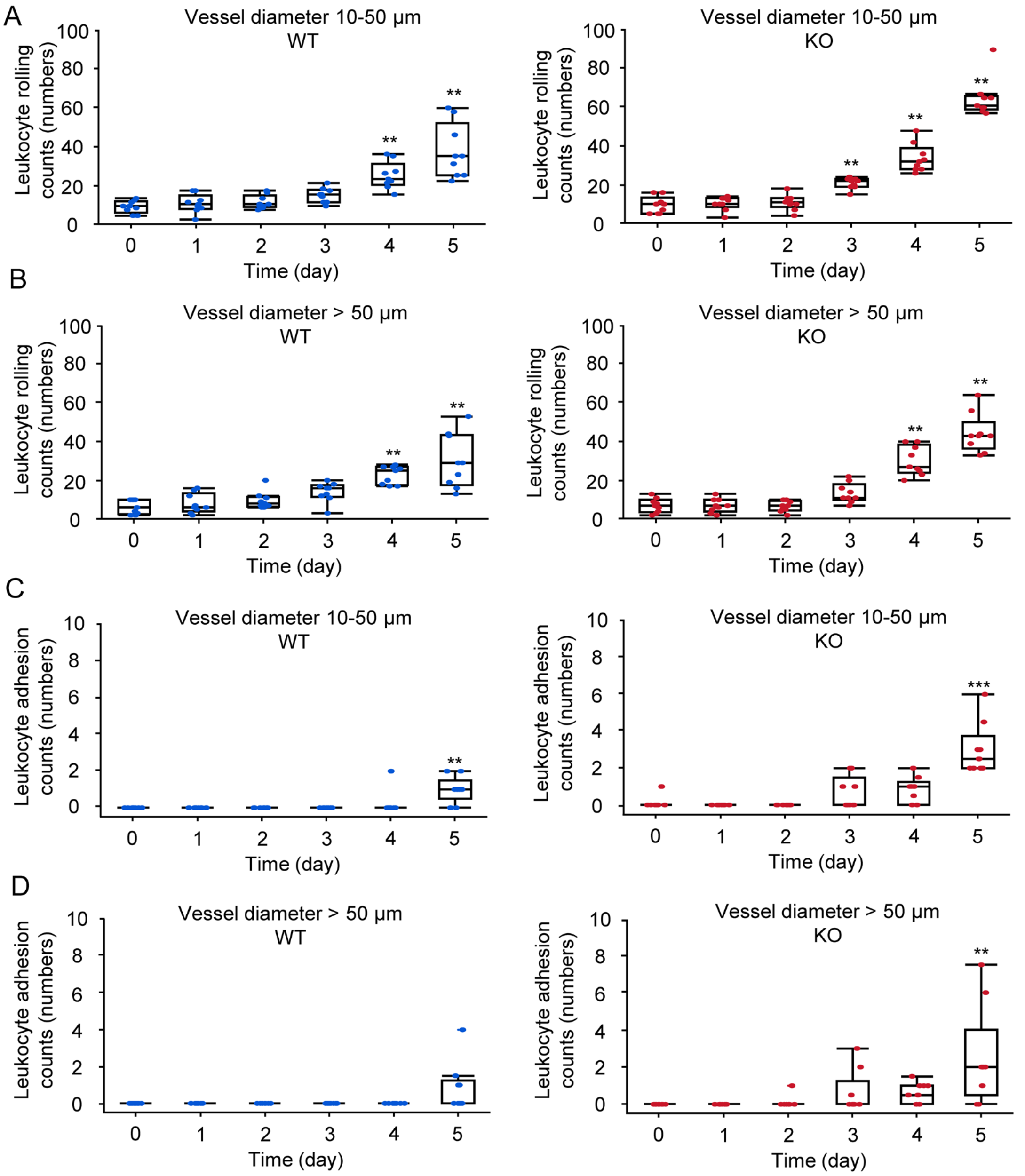


**Supplemental Figure 6. Details of leukocyte recruitment analysis**

Mice (age: 7–8 weeks, male, n=9 per group) were administered 5% DSS for 5 days. Leukocyte rolling in (A) 10–50 μm vessel and (B) vessel larger than 50 μm. Leukocyte adhesion in (C) 10–50 μm vessel and (D) vessel larger than 50 μm. Data are shown as median (centerline) and

25th and 75th percentiles (box bounds), with whiskers (maximum and minimum data points). Statistical significance was assessed by comparing the data from Day 0 with the those from other days using Steel's multiple comparison test. Blue dots indicate WT mice; red dots indicate ADAMTS13-KO mice. ** $p<0.01$; *** $p<0.001$.

### Supplemental Figure 7

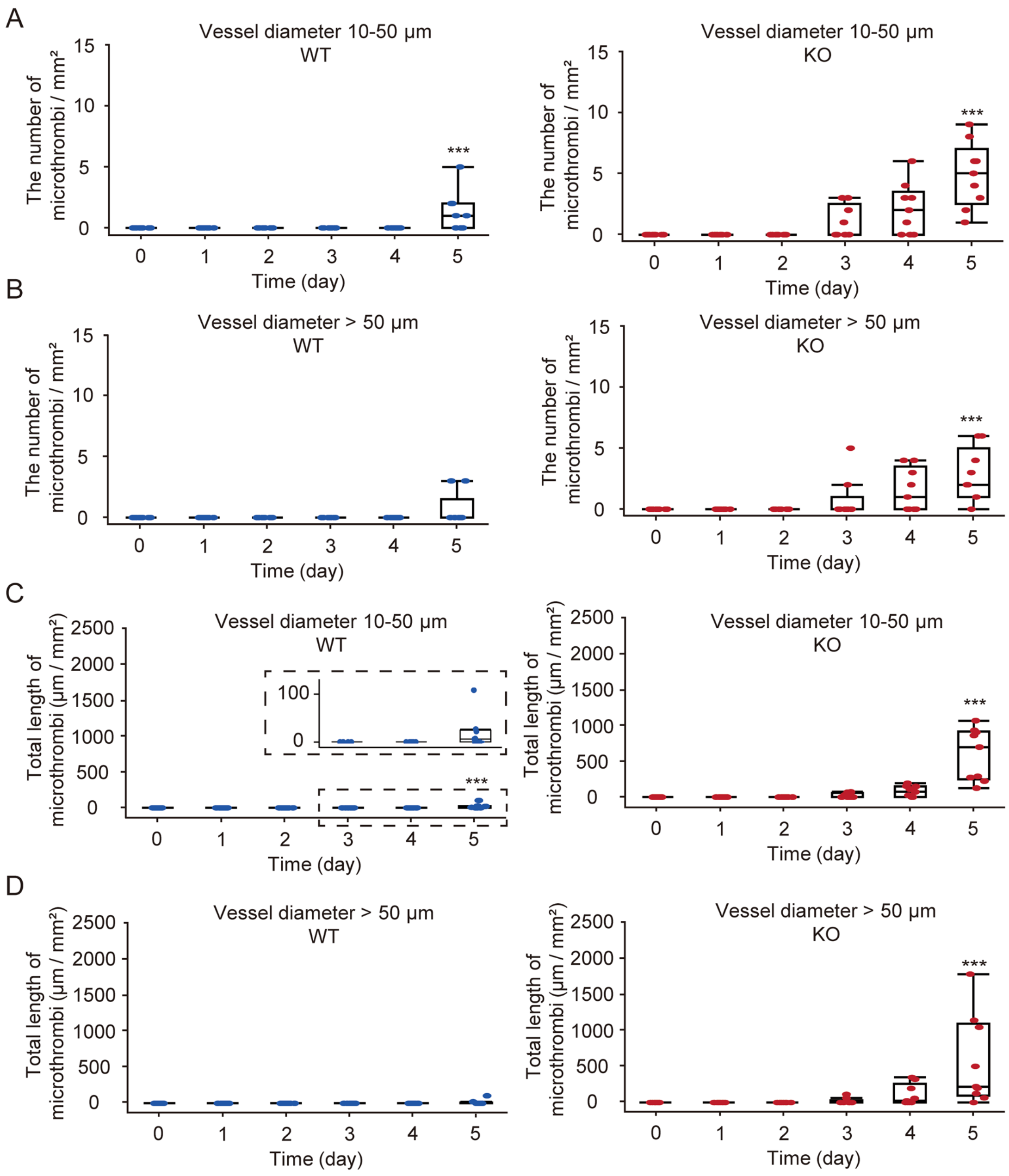


**Supplemental Figure 7. Details of microthrombi analysis**

Mice (age: 7–8 weeks, male, n=9 per group) were administered 5% DSS for 5 days. The number of microthrombi in a large observation area (1 × 1 mm field) of (A) 10–50 µm vessel and (B) vessel larger than 50 µm. Total length of microthrombi in a large observation area of

(C) 10–50 μm vessel and (D) vessel larger than 50 μm. Data are shown as median (centerline) and 25th and 75th percentiles (box bounds), with whiskers (maximum and minimum data points). Statistical significance was assessed by comparing data from Day 0 with those of the other days using Steel's multiple comparison test. Blue dots indicate WT mice; red dots indicate ADAMTS13-KO mice. ***$p$<0.001.

### Supplemental Figure 8

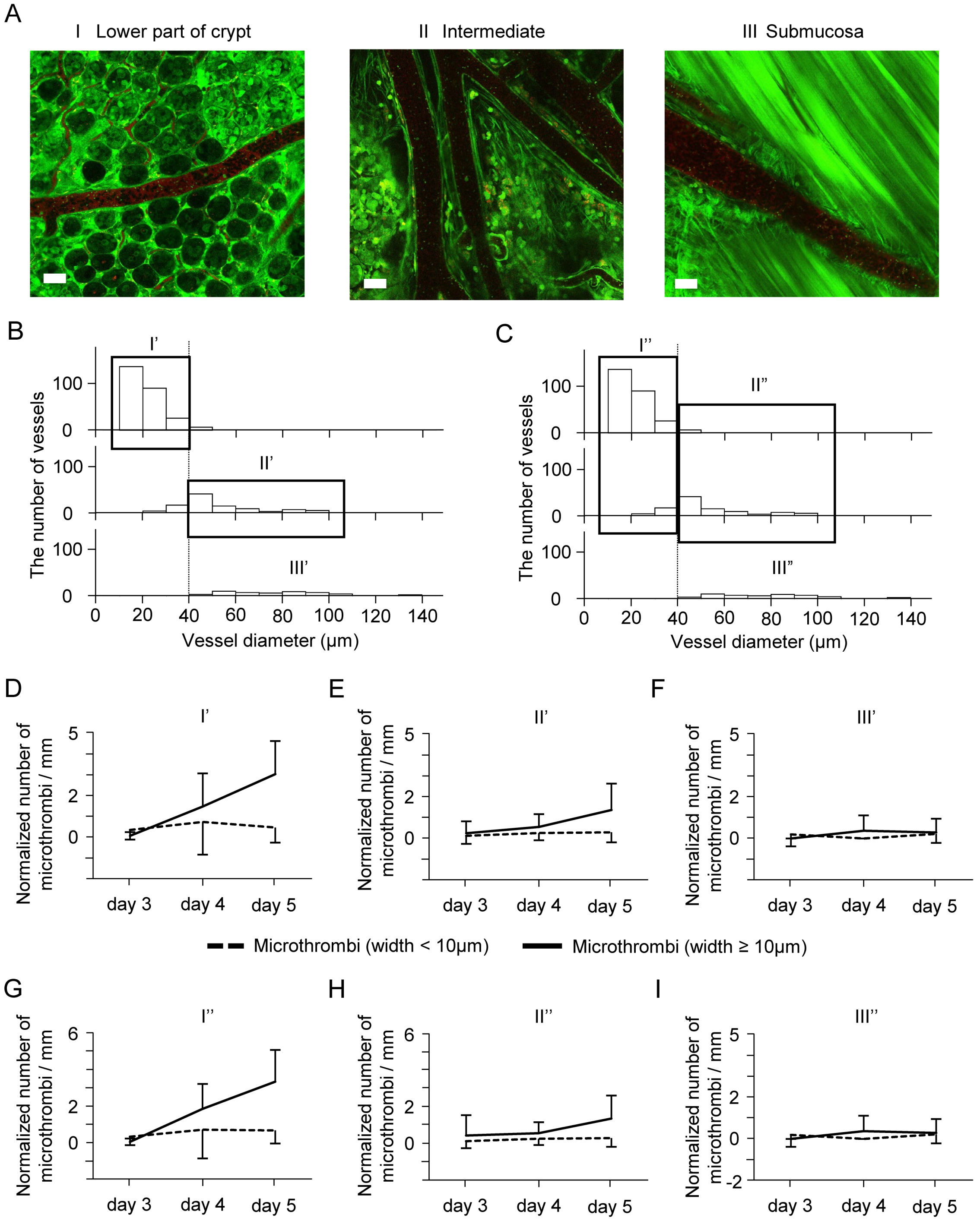


**Supplemental Figure 8. Sensitive spatiotemporal analysis of microthrombi**

ADAMTS13-KO Mice (age: 7–8 weeks, male, n=9) were administered 5% DSS for 5 days, and all observed microthrombi were analyzed. (A) Representative image of blood vessels categorized by anatomical location of colonic crypts. (B) Classification that excluded blood vessels with a diameter of 40 μm or more in Area I, and those with a diameter of less than 40 μm in Area II. (C) Classification that added blood vessels with a diameter of less than 40 μm from Area II to Area I, and those with a diameter of 40 μm or more from Area I to Area II. (D-F) Comparison of temporal changes in the normalized number of microthrombi at the 10 μm width boundary in vessels from I' (D), II' (E), and III' (F). The number of microthrombi per millimeter was calculated by normalizing to the length of the blood vessel. Microthrombi with a width of ≥ 10 μm at I' (D, black line) showed a significant increase over time ($p<0.001$). Microthrombi with the width < 10μm: dotted black line; microthrombi with the width ≥ 10μm: black line. (G-I) Comparison of temporal changes in the normalized number of microthrombi at the 10 μm width boundary in vessels from I'' (G), II'' (H), and III'' (I). Microthrombi with a width of ≥ 10 μm at I'' (G, black line) showed a significant increase over time ($p<0.001$). Data (D-I; mean ± SD) were analyzed by Dunnett's multiple comparison test to compare values recorded on day 3 with those on the other days.

**Supplemental Figure 9**

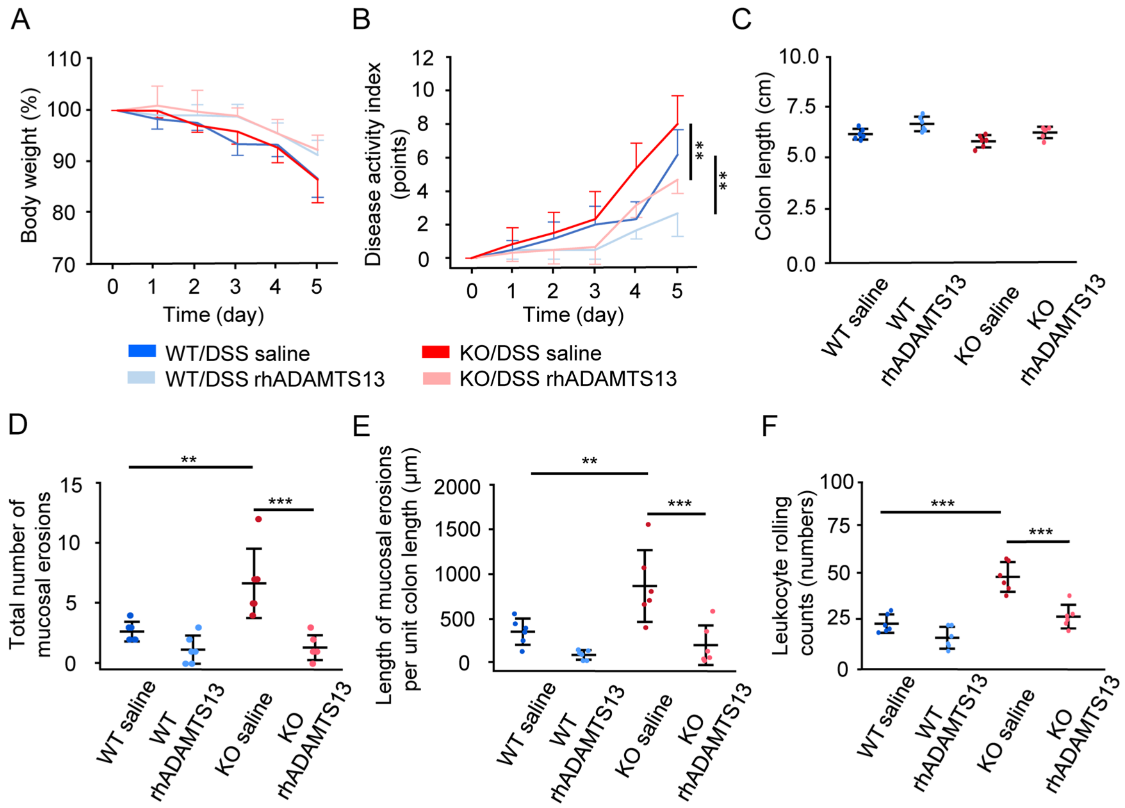


**Supplemental Figure 9**. Improvement of DSS-induced colitis after rhADAMTS13 treatment

Mice (age: 7–8 weeks, male, n=6 per group) were administered 5% DSS for 5 days and simultaneously received rhADAMTS13 (WT, light blue lines/dots; KO, light red lines/dots) or saline (WT, blue lines/dots; KO, red lines/dots) injected intravenously on days 1–5. (A) Body-weight changes are shown as percentages on Day 0. (B) The disease activity index is shown at all points. (C) Total colon length on Day 5. (D) Number of mucosal erosions in the total colon on Day 5. (E) Length of mucosal erosion per 1 cm of the colon on Day 5. (F) Leukocyte rolling count on Day 5. Data (mean ± SD) were analyzed by the Student's t-test with Bonferroni correction. **p<0.01; ***p<0.001.

**Supplemental Video legends**

**Supplemental Video 1**. Representative 3D image of colonic crypt in ADAMTS13-KO mice without colitis after TRITC-dextran administration

IMARIS 8 (v8.1, Bitplane, Belfast, Northern Ireland, UK) was used to create the 3D images. GFP (green), TRITC (red).

**Supplemental Video 2**. Representative 3D image of colonic crypt in ADAMTS13-KO mice with colitis after TRITC-dextran administration

IMARIS 8 (v8.1, Bitplane, Belfast, Northern Ireland, UK) was used to create the 3D images. GFP (green), TRITC (red).

**Supplemental Video 3**. Representative movie of leukocyte rolling and adhesion in 10–50 μm blood vessels of WT mice treated with DSS on Day 0.

Recording duration: 1 min. Recording frame rate: 7 s/frame. Data correspond to the images in Figure S5B, S5C, and S5E; Figure S6A and S6C. Scale bar: 50 μm.

**Supplemental Video 4.** Representative movie of leukocyte rolling and adhesion in 10–50 μm blood vessels of ADAMTS13-KO mice treated with DSS on Day 0.

Recording duration: 1 min. Recording frame rate: 7 s/frame. Data correspond to Figure S5B, S5C, and S5E; Figure S6A and S6C. Scale bar: 50 μm.

**Supplemental Video 5**. Representative movie of leukocyte rolling and adhesion in 10–50 μm blood vessels of WT mice treated with DSS for 5 days.

Recording duration: 1 min. Recording frame rate: 7 s/frame. Data correspond to Figure S5B, S5C, and S5E; Figure S6A and S6C. Scale bar: 50 μm.

**Supplemental Video 6**. Representative movie of leukocyte rolling and adhesion in 10–50 μm blood vessels of ADAMTS13-KO mice treated with DSS for 5 days.

Recording duration: 1 min. Recording frame rate: 7 s/frame. Data correspond to Figure S5B, S5C, and S5E; Figure S6A and S6C. Scale bar: 50 μm.

**Supplemental Video 7**. Representative movie of microvascular thrombi in 10–50 μm blood vessels of WT mice treated with DSS for 5 days.

GFP (green) and anti-VWF antibodies (red). Recording duration: 1 min. Recording frame rate: 7 s/frame. Data correspond to Figure 4C, 4D, 4G, and 4H; Figure S7A and S7C. Scale bar: 50 μm.

**Supplemental Video 8**. Representative movie of microvascular thrombi in 10–50 μm blood vessels of ADAMTS13-KO mice treated with DSS for 5 days.

GFP (green) and anti-VWF antibodies (red). Recording duration: 1 min. Recording frame rate: 7 s/frame. Data correspond to Figure 4E through 4H; Figure S7A and S7C. Scale bar: 50 μm.
